# Adult born hippocampal granule cells promote pattern separation by bidirectionally modulating the remapping of place and cue cells

**DOI:** 10.1101/2022.12.08.519632

**Authors:** Sebnem N. Tuncdemir, Andres D. Grosmark, Hannah Chung, Victor M. Luna, Clay O. Lacefield, Attila Losonczy, Rene Hen

## Abstract

The hippocampal dentate gyrus (DG) exhibits a unique form of neural plasticity that results from continuous integration of adult born neurons, referred to as ‘adult neurogenesis’. Recent studies have proposed that adult neurogenesis promotes the ability to encode new memories without interference from previously stored memories that share similar features, through a neural computation known as pattern separation. However, due to lack of *in vivo* physiological evidence, the manner in which adult neurogenesis contributes to pattern separation remains unknown. Here, we investigate the contribution of functionally integrated yet immature adult born granule cells (iGCs) to DG computations by examining how chronic ablation or acute chemogenetic silencing of iGCs affects the activity of mature granule cells (mGCs) using *in vivo* 2-photon Ca^2+^ imaging. In both cases we observed altered remapping of mGCs but in opposite directions depending on their tuning selectivity. Rather than broadly modulating the activity of all mGCs, iGCs promote the remapping of place cells but limit the remapping of mGCs representing sensory cues (cue cells). We propose that these properties of iGCs explain their role in pattern separation because they promote the formation of non-overlapping representations for identical sensory cues encountered in different locations. Conversely, the absence of iGCs shifts the DG network to a state dominated by sensory cue information, a situation that is consistent with the overgeneralization often observed in anxiety disorders such as PTSD.

## Introduction

The dentate gyrus (DG) of the hippocampus is one of two areas in the adult mammalian brain in which new neurons are continuously generated and integrated into the existing local DG circuit over several weeks. Four to 6 weeks after their birth, immature granule cells (iGCs) exhibit increased synaptic plasticity and intrinsic excitability compared to mature granule cells (mGCs, >8 weeks old) (Snyder et al., 2001; Schmidt-Hieber et al., 2004; Ge et al., 2007; Marín-Burgin et al., 2012). Acute or chronic manipulations of iGCs during this period have demonstrated an important role for iGCs in hippocampus-dependent tasks involving behavioral pattern separation, such as the ability to discriminate between similar contexts or spatial locations (Sahay et al., 2011; Denny et al., 2012; Gu et al., 2012; Nakashiba et al., 2012; Danielson et al., 2016; Huckleberry et al., 2018). Notably, the deficits resulting from impaired hippocampal neurogenesis are consistent with the overgeneralization often associated with mood and anxiety disorders, specifically post-traumatic stress disorder (Lissek et al., 2010; Kheirbek et al., 2012; Belzung et al., 2015; Lange et al., 2017). It has therefore been suggested that iGCs play a role in this common endophenotype of neuropsychiatric disorders.

The behavioral impact of iGCs has been proposed to result from their inhibitory modulation of mGCs resulting in increased sparseness of DG representations (Lacefield et al., 2012; Ikrar et al., 2013; Drew et al., 2016; Anacker et al., 2018; McHugh et al., 2022). However, rather than broadly modulating mGCs in a uniform fashion, iGCs appear to dynamically interact with mGCs such that under different cortical inputs as engaged by distinct cognitive demands, iGCs may either inhibit or excite mGCs (Luna et al., 2019). Thus, iGCs may engage different modulatory pathways to promote the formation of distinct mGC cell assemblies representing distinct contexts without interference from previously stored contextual memories.

A large proportion of principal neurons in all subfields of the hippocampus, including those in the DG, are considered place cells because they fire at specific locations corresponding to the position of an animal within an environment through the integration of location-related sensory and self-motion information (O’Keefe and Dostrovsky, 1971; Burgess and O’Keefe, 1996). Place cells change their preferred firing location and their firing rate when the environment changes, a process known as ‘remapping’ (Latuske et al., 2017; Kubie et al., 2020). Remapping of place cells is thought to support spatial and contextual discrimination by enforcing unique representations of distinct environments and behavioral experiences with varying levels of overlap with previously stored ones (Colgin et al., 2008). While remapping has been found in all hippocampal subregions, this feature is most consistent with the proposed pattern separation function of the DG (Yassa and Stark, 2011), where similar inputs are represented by non-overlapping populations resulting in increased selectivity to similar contexts (Marr, 1971; Knierim and Neunuebel, 2016). While mGCs have been shown to exhibit global or rate remapping (Diamantaki et al., 2016; Danielson et al., 2017; GoodSmith et al., 2017; Senzai and Buzsáki, 2017; Hainmueller and Bartos, 2018), how iGCs influence the remapping of mGCs in different contexts is not known. In addition, recent work has begun to characterize a functional diversity within the mGC population, whose activity are modulated by space or sensory cues (place cells and cue cells) (Jung et al., 2019; Woods et al., 2020; GoodSmith et al., 2022; Tuncdemir et al., 2022). These spatial and cue representations are distinct in their stability across different contexts as well as long-term dynamics (Tuncdemir et al., 2022), suggesting that different subpopulations of mGCs encode distinct contextual features. But, whether iGCs have a differential effect on the sensory cue versus place encoding mGCs or their remapping dynamics is not known.

Here, we have investigated the contribution of iGCs to DG computations, by examining how chronically ablating or chemogenetically silencing iGCs affects the activity of mGCs using *in vivo* 2-photon Ca^2+^ imaging. We find that chronically ablating or transiently inhibiting iGCs resulted in decreased remapping of mGCs spatial representations when presented with different contexts. However, rather than broadly modulating Ca^2+^ firing rates or spatial tuning of all mGCs, iGCs appear to modulate mGCs bidirectionally depending on their tuning specificity: they impede the remapping of mGCs representing sensory cues (cue cells) but promote the remapping of place cells.

## Results

### Immature granule cells are necessary for the remapping of spatial representations

To investigate how iGCs influence DG network activity, we first used a well-established method of bilateral X-irradiation to ablate adult born immature GCs (Santarelli et al., 2003; Burghardt et al., 2012) and recorded Ca^2+^ activity in large populations of mGCs in the mouse dorsal DG during head-fixed locomotion on a treadmill (Danielson et al., 2016). After 2 months of recovery period after irradiation, mice were injected with a recombinant adeno-associated virus (rAAV) expressing the fluorescent calcium indicator GCaMP6s, implanted with a chronic imaging window over the dorsal DG, habituated to head-fixation and trained to run on a featureless treadmill belt in order to receive randomly delivered operant water rewards (Figure 1A). Ablation of neurogenesis was confirmed by the complete loss of doublecortin immunoreactive cells in the DG of X-irradiated mice (X-IR) compared to controls (Figure 1B). To study how the loss of adult hippocampal neurogenesis affects the specificity of spatial representations within the mGC population, mice were recorded during three consecutive exposures to two identical and then a different context. These contexts consisted of three fabric materials joined together, and contained tactile cues fixed in the middle of each section together with olfactory, visual and auditory cues delivered ambiently inside the recording rig (Figure 1A, (Danielson et al., 2016)). The contexts A and B had the same treadmill belts and affixed tactile cues that were reordered along with different ambient olfactory, visual and auditory cues. We tracked the same cells over three consecutive sessions (Sheintuch et al., 2017; Zaremba et al., 2017) and were able to cross-register substantial numbers of the GCs active in the same fields of view in both control and X-IR mice during sequential exposures to either same (A-A’) or different (B) contexts (Figures S1A-E).

**Figure 1:**
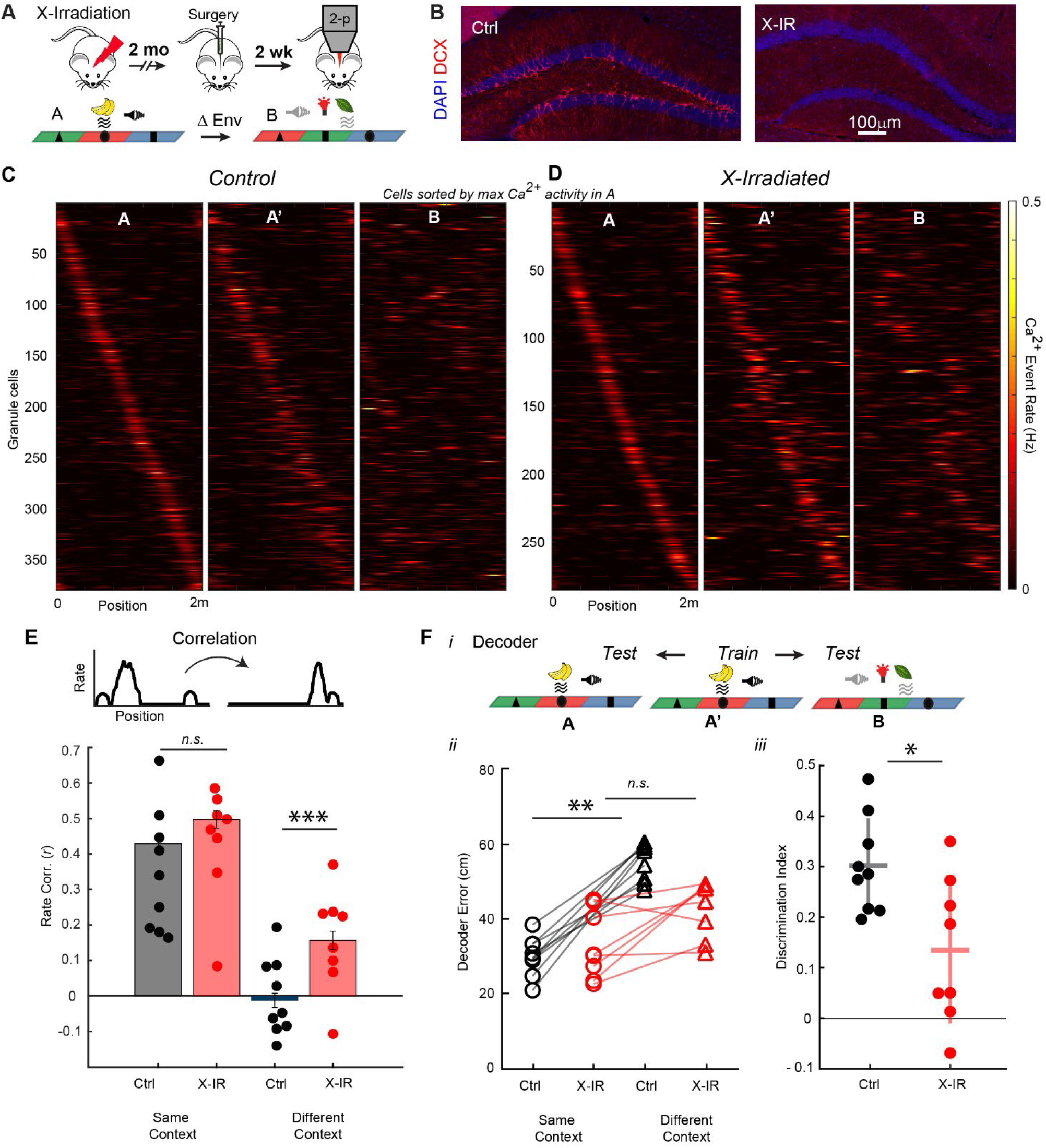
Chronic ablation of neurogenesis impairs context discrimination in the dentate gyrus. **A)** (Top) Adult hippocampal neurogenesis was arrested using focal X-irradiation (X-IR) of the hippocampus bilaterally, followed by virus injection and window implantation 2 months later. Head-fixed mice were subsequently trained to run on a featureless belt to find randomly scattered uncued rewards. (Bottom) Mice were then tested first in context A for 15 min, re-exposed to context A for 15 min, and then tested in context B for 15 min with 1 hr between runs. Contexts A and B consist of distinct ambient odor, tone and light stimuli but the same three joined fabrics (total length of 2m) with different textures and objects that are shuffled between contexts A and B. **B)** Doublecortin expressing adult born immature granule cells (iGCs, red) are completely eliminated in X-IR mice (right). Blue is DAPI. (Scale bar = 100mm). **C)** Spatial firing rate maps during 3 sequential exposures to the same and different contexts in control mice. The second exposure to Context A is shown as A’. Cells were selected based on having a place field in at least one session and are arranged based on maximum events rate in context A. Each row across all graphs represents a single cell matched across all contexts. N_Ctrl_=380 tuned cells from 9 mice. **D)** Spatial firing rate maps during 3 sequential exposures to the same and different contexts in X-IR mice. N_X-IR_= 285 tuned cells from 8 mice. **E)** Top: Similarity of spatial representations of cells with a field in at least one context were measured by calculating Pearson’s correlations of rate maps between sequential exposures to the same (A-A’) or different (A’-B) contexts (top). Bottom: Bar plots represent group data; circles represent per-mouse data. Interaction *F*_1,30_ = 10.89, \*\**P*=0.0025; Context *F*_1,30_ = 368.6, \*\*\**P*<0.0001; Genotype *F*_1,30_ = 47.93, \*\**P*<0.0001; 2-way ANOVA. Same context: *P*_*Ctrl* – *IR*_ = 0.0705; Different context: \*\*\**P*_*Ctrl* – *IR*_ <0.0001; Tukey’s multiple comparisons test. ***F)*** Position decoded by a Bayesian algorithm among granule cells found in all sessions by training using neural activity in A’ and calculating predicted position in the same (A) or different (B) contexts (i). Line graphs represent median spatial decoder error calculated as the difference between the predicted position in the training data from those in test datasets that are higher in control animals compared to X-IR when spatial activity from different environments was used to decode position (ii). Control: *W* = 45, \*\**P* _*Same Ctx* – *Different Ctx*_ = 0.0039; X-IR: *W* = 28, *P* _*Same Ctx* – *Different Ctx*_ = 0.0604; Wilcoxon rank sum test. Discrimination index was calculated as: (Error _Different Context_ - Error _Same Context_)/ (Error _Different Context_ + Error _Same Context_), *U*=13, \**P*_*Ctrl* – *IR*_ *= 0*.*0*274, Mann-Whitney test. Control: 97.14 ± 13.86, X-IR: 93.57 ± 9.69 granule cells matched in test and train sessions. Error bars, ± sem (iii, *see also Supplementary Figure 1)*

We focused on the mGCs that showed spatially tuned activity in at least one of the three sessions and compared the similarity of mGCs’ spatial fields between two sequential exposures to either same (A-A’) or different (A’-B) contexts. Both groups of mice displayed stable spatial maps when they were exposed to the same context and distinct maps when exposed to different contexts (Figure 1C,D). Strikingly, X-IR mice showed markedly lower levels of reorganization of mGC spatial firing compared to controls in a different context (Figure 1C,D). Accordingly, spatially tuned mGCs displayed high correlation of their tuning curves in both controls and X-IR mice in the same context, whereas X-IR mice displayed significantly higher correlation of tuning curves compared to controls in different contexts (Figure 1E). In addition to responses of individual mGCs, we were also interested in examining population level changes in response to chronic ablation of neurogenesis. Hence, we calculated population level differences of spatial coding between the two groups of mice by using a Bayesian decoder (Grosmark et al., 2021) to predict the position of the mice in context A’, by training the decoder with the firing rate vectors of mGCs, found in the same (A) or different (B) contexts (Figure 1Fi), regardless of their spatial tuning specificity. While the predicted position error was similar in both groups of mice when position was predicted using the cells found in the same environment (A-A’), the X-IR mice showed consistently lower prediction errors compared to controls in different contexts (A’-B, Figure 1Fii) suggesting impaired context discrimination in X-IR mice compared to controls (Figure 1Fiii).

We further confirmed that the differences in tuning correlation or decoder prediction error in different contexts between control and X-IR mice did not result from the differential cross-session registration of cells due to differences in the stability of fields of view. There were no differences in the fraction of cells cross-registered in all three consecutive sessions between control and X-IR mice, as assessed by the normalized fraction of all active or only spatially tuned mGCs in the field of view (Figure S1C). Similarly, the fraction of all or only spatially tuned mGCs cross-registered in contexts A or B were comparable between different groups (Figures S1D,E). Additionally, the population vector correlations of the Ca^2+^ event rates of mGCs in X-IR mice at all positions in different contexts (A vs B) were increased compared to controls (Figures S1F,G), consistent with a role for adult neurogenesis in promoting the remapping of spatial representations during exploration of different contexts.

X-irradiation permanently blocks hippocampal neurogenesis; hence, we sought to acutely and selectively silence the activity of 4-week-old iGCs because this time point is within the hyperplastic phase during which they are thought to most strongly influence behavior (Denny et al., 2012; Gu et al., 2012). We expressed the inhibitory designer receptor exclusively activated by designer drugs (DREADD), *hM4Di*, in iGCs by crossing a transgenic mouse line with a tamoxifen-inducible CreERT2 recombinase, expressed under the control of the *Ascl1* gene promoter (Kim et al., 2011) to mice expressing STOP-floxed-*hM4Di* (Ray et al., 2011; Anacker et al., 2018) (*Ascl-creERT2*^+/−^;*loxP*-*stop-mCherry-loxP-hM4Di*, referred to as *Ascl*^*CreER*^ *Di* ^F/F^ compared to *Cre* negative *Di* ^F/F^ littermates). Upon binding of the DREADD receptor agonist clozapine-*N*-oxide (CNO), hM4Di activates the canonical Gi pathway resulting in hyperpolarization of 4-week-old iGCs. 2 weeks after tamoxifen injection, *Ascl*^*CreER*^ *Di* ^F/F^ mice and *Di* ^F/F^ controls were implanted with a chronic imaging window over the dorsal DG and were injected with rAAV expressing the fluorescent calcium indicator GCaMP7s. Over the following 2 weeks, mice were habituated to head-fixation and trained to run on a treadmill belt; next, we investigated the spatial activity in mGCs after acute inhibition of 4-week-old iGCs (Figure 2A). The efficacy of this strategy was further confirmed in acute slice preparations (Figure S2A). In a separate cohort of *Ascl*^*CreER*^ mice crossed to a STOP-floxed *TdTomato* line (Madisen et al., 2010), we confirmed that this experimental timeline and the titer of rAAV injection did not result in a significantly reduced number of iGCs in the injected hemispheres compared to the contralateral hemisphere (Figures 2B, S2B, (Johnston et al., 2021)).

**Figure 2:**
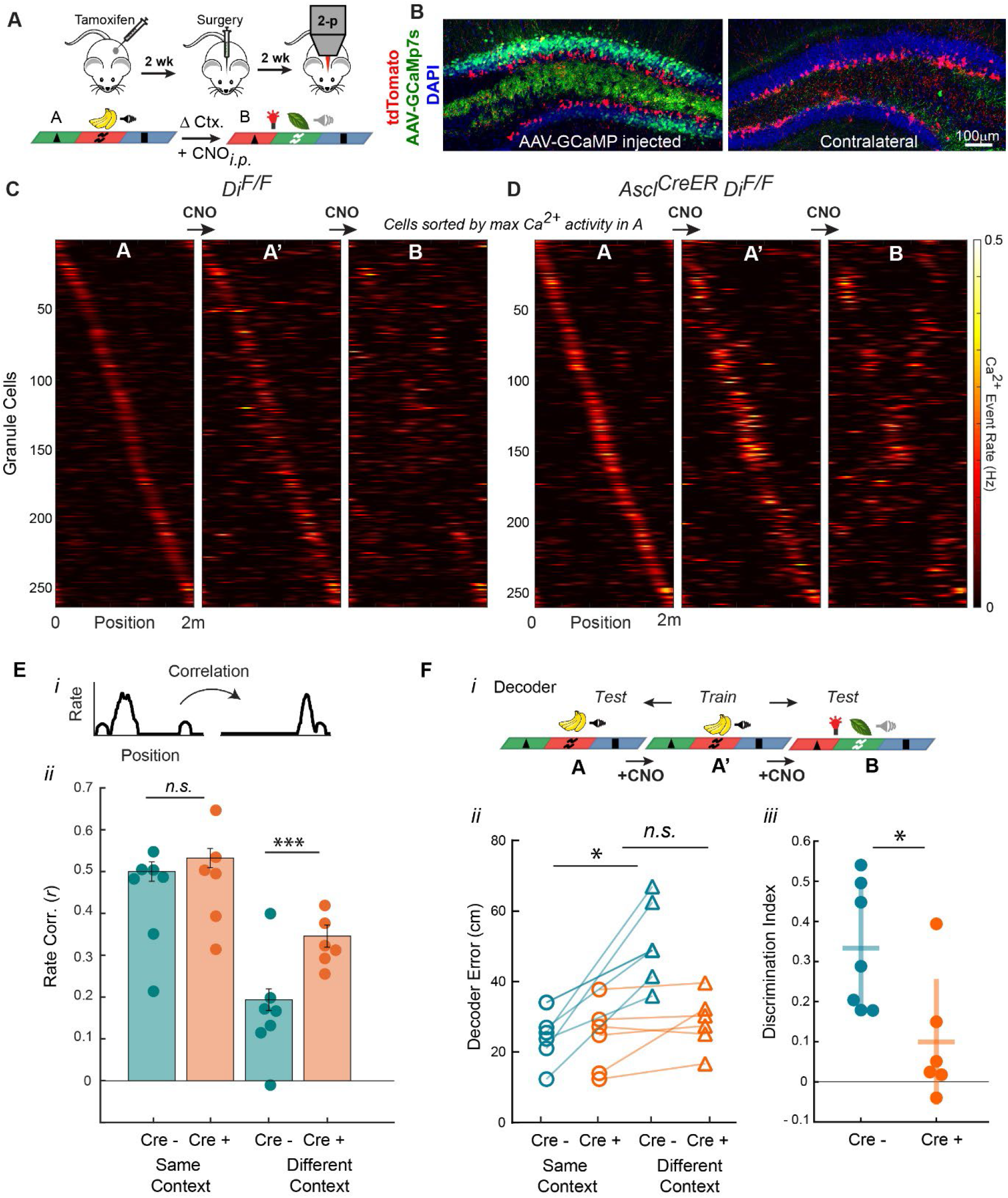
Acute silencing of 4-week-old immature granule cells impairs context discrimination in the DG. **A)** Adult born immature granule cells were silenced using designer receptor exclusively activated by designer drugs (hM4Di), which activates inhibitory Gi-signaling upon stimulation by the ligand, clozapine-N-oxide (CNO). Tamoxifen (TAM) inducible Ascl^CreER^;Di^F/F^ mice and Di^F/F^ controls were injected with Tamoxifen (100mg/kg) for 5 consecutive days, and followed by rAAV injection and window implantation 2 weeks later. Head-fixed mice were subsequently trained to run on a featureless belt to find randomly scattered hidden rewards. (Bottom) Mice were tested in different Contexts as in Figure 1 but different olfactory stimuli were delivered in the middle of one of the belt sections (1sec.). First session in context A was imaged as a baseline, then all mice received CNO (5mg/kg) *i*.*p*. injections 30 min before running in contexts A’ and B while at home cage. **B)** A separate cohort of Ascl^CreER^;TdTomato^F/F^ mice were injected with TAM followed by AAV-GCaMp7s injection 2 weeks later. AAV injection did not result in differences in genetically targeted iGCs in the injected hemisphere compared to the contralateral hemisphere, for quantification see Figure S2A. **C)** Spatial firing patterns during 3 sequential exposures to the same and different contexts in Di^F/F^ control mice. The second exposure to Context A is shown as A’. Cells selected based on having a place field in at least one session and are arranged based on maximum events rate in A. Each row across all graphs represents a single cell matched across all contexts. N_Cre-_=261 cells, 7 mice. **D)** Spatial firing patterns during 3 sequential exposures to the same and different contexts in Ascl^CreER^;Di^F/F^ mice. N_Cre+_= 275 cells, 6 mice. **E)** Top: Similarity of spatial representations of cells with a field in at least one context are measured by calculating Pearson’s correlations of rate maps between sequential exposures to the same (A-A’) or different (A’-B) contexts (top). Bottom: Bar plots represent group data; circles represent per animal data. Interaction *F*_1,22_ = 11.15, \*\**P*=0.0030; Context *F*_1,22_ = 191.9, \*\*\**P*<0.0001; Genotype *F*_1,22_ = 34.89, \*\**P*<0.0001; 2-way ANOVA. Same context: *P*_*Cre-* – *Cre*+_ = 0.2928; Different context: \*\*\**P*_*Cre-* – *Cre*+_ <0.0001; Tukey’s multiple comparisons test. **F)** Position decoded by a Bayesian algorithm among cells found in all sessions by training using neural activity in A’ and calculating predicted position in the same (A) or different (B) contexts (i). Line graphs represent median spatial decoder error calculated as the difference between the predicted position in the training data from those in test datasets that is higher in Cre negative Di^F/F^ animals compared to Ascl^CreER^;Di^F/F^ when spatial activity from different environments was used to decode position (ii). Di^F/F^: *W* = 28, \**P* ^*Same Ctx – Different Ctx*^ = 0.0156; Ascl^CreER^;Di^F/F^: *W* = 15, *P* ^*Same Ctx – Different Ctx*^ = 0.1563; Wilcoxon rank sum test. iii. Discrimination index was calculated as: (Error _Different Context_ - Error _Same Context_)/ (Error _Different Context_ + Error _Same Context_), *U*=4, \**P*_*Cre—Cre+*_ *= 0*.*0*140, Mann-Whitney test. Di^F/F^: 83.42 ± 5.89, Ascl^CreER^;Di^F/F^: 83 ± 8.68 granule cells matched in test and train sessions. Error bars, ± sem. (iii, *see also Supplementary Figure 2)*

Like in the previous experiment, we compared neural activity during two sequential exposures of either identical or different contexts. The first session in context A was the baseline, and CNO was *i*.*p* injected prior to re-exposure to A (A’) and also before exposure to the different context B in both groups of mice (Figure 2A, Methods). The injection of CNO did not have an effect on the spatial firing stability in the same contexts in both groups of mice (Figure 2C,D). However, acute inhibition of 4-week-old iGCs in *Ascl*^*CreER*^ *Di* ^F/F^ mice resulted in impaired remapping in different contexts (Figure 2C,D). Similar to the differences between controls and X-IR mice: *Cre*-negative *Di* ^F/F^ controls did remap their firing fields between contexts A and B whereas this ability was reduced in *Ascl*^*CreER*^ *Di* ^F/F^ mice. mGCs with spatially tuned activity in at least one of the contexts A, A’ or B (same selection criteria as above) displayed high correlation of their tuning curves in both *Di* ^F/F^ controls and *Ascl*^*CreER*^ *Di* ^F/F^ mice in the same context (A-A’), whereas *Ascl*^*CreER*^ *Di* ^F/F^ mice displayed significantly higher correlation of tuning curves compared to controls in different contexts (A’-B, Figure 2E). Bayesian decoding of spatial locations also reflected remapping impairments in CNO-treated *Ascl*^*CreER*^ *Di* ^F/F^ mice compared to CNO-treated controls (Figures 2Fi,ii), resulting in significantly lower context discrimination in *Ascl*^*CreER*^ *Di* ^F/F^ mice compared to *Di* ^F/F^ controls (Figure 2Fiii). Consistent with the chronic ablation experiments, the correlation of the mGCs in *Ascl*^*CreER*^ *Di* ^F/F^ mice at all positions of the different contexts (A vs B) were increased compared to *Di* ^F/F^ controls (Figures S2C,D). Further, the effect size of Control - X-IR and *Di* ^F/F^ - *Ascl*^*CreER*^ *Di* ^F/F^ cohorts were comparable (Figure S2E), indicating that the observed impairments in both irradiation- and chemogenetic-based mouse models likely result from cell autonomous effects of iGCs on the DG circuitry rather than non-specific effects of irradiation (Hill et al., 2015). Hence, our results with chronic ablation of adult hippocampal neurogenesis and acute silencing of iGCs reveal that 4-week-old iGCs facilitate neural discrimination of different contexts by promoting the remapping of mGCs’ spatial receptive fields.

### Immature granule cells modulate remapping without changing global activity levels or spatial tuning

Next, we examined the mechanisms by which iGCs promote spatial coding flexibility. Numerous studies have proposed that iGCs enforce sparse and distributed activity within the mGC population via lateral inhibition or competition with excitatory synapses (Lacefield et al., 2012; Anacker et al., 2018; McHugh et al., 2022). Yet, we recently demonstrated that depending on the incoming afferents from the lateral or medial entorhinal cortex, iGCs can either inhibit or excite mGCs rather than uniformly inhibiting all mGCs (Luna et al., 2019). Here, we found that the fraction of total active (with at least 1 Ca^2+^ transient during 15 min recording) and spatially tuned mGCs did not significantly change after chronic ablation (Figure 3A) or acute inhibition of iGCs (Figure 3D). In both models, there were no significant differences in overall mean Ca^2+^ event rates within a session per animal (Figures 3B, E) and spatially tuned neurons had comparable spatial information content (Skaggs et al., 1993) (Figures S3 A,D). In addition, similar fractions of neurons across genotypes imaged during Context A’ maintained (Figures S3B,E) or lost (Figures S3C,F) their spatial tuning when cross-session registered to cells in same or different contexts in both mouse models. Instead, the relationship of individual mGC’s Ca^2+^ event rates with their remapping index, a measure of the difference in correlations between similar contexts and different contexts (Senzai and Buzsáki, 2017), were consistently different from controls in both models of chronic X-IR ablation or acute DREADD inhibition of iGCs. The distribution of the Ca^2+^ event rates of spatially tuned mGCs revealed that the high firing rate cells in X-IR mice showed the most pronounced deficit in context selectivity (as defined by their low remapping index) compared to controls (Figure 3C). Moreover, we tracked how transient inhibition of iGCs affects both Ca^2+^ event rates (Figure S3G) and the remapping responses (Figure S3H) of single neurons before and after injection of CNO. We show that mGCs most sensitive to the effect of iGCs in *Ascl*^*CreER*^ *Di* ^F/F^ mice were those that were disinhibited and exhibited the least remapping between contexts A-B compared to controls (Figure 3F). Taken together, these results demonstrate that chronic ablation and acute inhibition of iGCs do not influence every mGC evenly, suggesting a functional heterogeneity in the way iGCs engage different modulatory pathways to promote the formation of distinct spatial maps by mGCs. Our results also reveal that under normal conditions, the subpopulation of mGCs that are selectively inhibited by iGCs display increased context selectivity.

**Figure 3:**
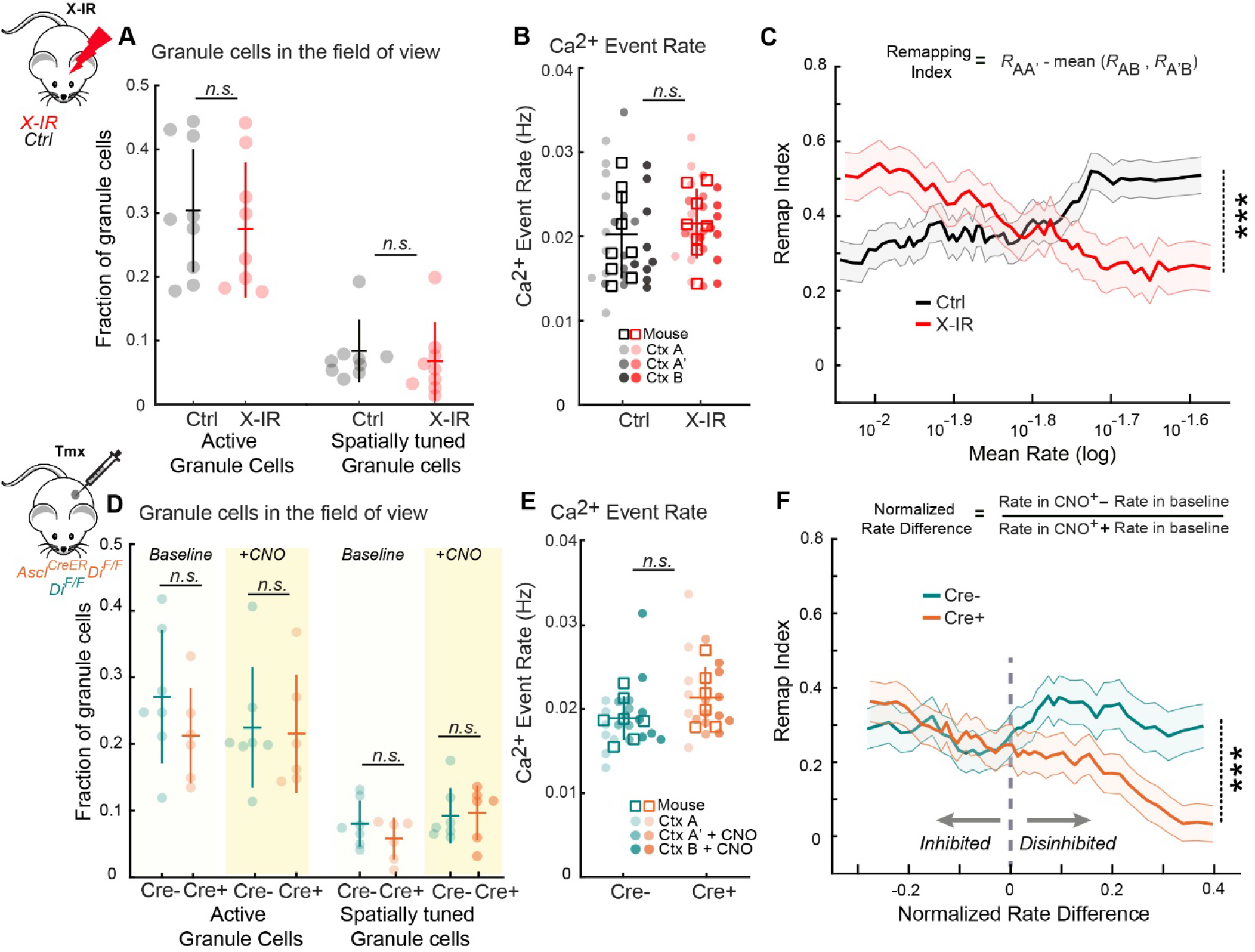
Immature granule cells modulate stability of representations without changing global activity levels or spatial selectivity. **A)** Fraction of active granule cells (GC, with at least 0.001 transients per s) and spatially tuned cells in the imaging field of view during context discrimination task. Active granule cells: *U*=23, *P*_*Ctrl—IR*_ *= 0*.*23*, spatially tuned cells: *U*=26, *P*_*Ctrl—IR*_ *= 0*.*37*, Mann-Whitney test. Control: 166.22 ± 7.48, 40.92 ± 4.36, X-IR: 144.54 ± 6.36, 30.04 ± 3.82 granule cells and spatially tuned cells, respectively. **B)** Spatial Ca^2+^ event rates from during running bouts. Squares represent per mouse circles represent per session averages of spatial firing rates. Interaction *F*_2,45_ = 0.03, *P*=0.96; Context *F*_2,45_ = 0.45, *P*=0.63; Genotype *F*_1,45_ = 0.65, *P*=0.42; 2-way ANOVA. **C)** Granule cells’ averaged Ca^2+^ rates are plotted against their remapping index in control and X-IR mice within a sliding 25 percentile-wide window of the averaged rates on a log scale. Context selectivity was calculated by subtracting the mean of the AB and A’B correlations from the AA’ correlation GCs with higher mean rates have significantly lower remapping index in X-IR mice compared to those in controls. Pearson’s *R*_Ctrl_=0.1628, *R*_IR_=-0.1405, \*\*\**P= 0*.*000340*, Fisher Z test. **D)** Fraction of active granule cells and spatially tuned cells in the imaging field of view before and after CNO injection in the second exposure to context A in *Cre* negative Di^F/F^ controls (green) compared to Ascl^CreER^;Di^F/F^ mice (orange). Active granule cells: Interaction *F*_1,22_ = 0.49, *P*=0.48; CNO *F*_1,22_ = 0.38, *P*=0.54; Genotype *F*_1,22_ = 0.94, *P*=0.34; Spatially tuned cells: Interaction *F*_1,22_ = 1.06, *P*=0.31; CNO *F*_1,22_ = 3.43, *P*=0.078; Genotype *F*_1,22_ = 0.57, *P*=0.45; 2-way ANOVA. Di^F/F^: 141.76 ± 7.90, 45.52 ± 3.25, Ascl^CreER^;Di^F/F^: 135.55 ± 6.35, 52 ± 7.87 granule cells and spatially tuned cells, respectively. **E)** Spatial Ca^2+^ event rates from during running bouts (squares represent per mouse, circles represent per session averages of spatial firing rates). Darker shades represent sessions with CNO injections. Interaction *F*_2,33_ = 0.001, *P*=0.99; Context *F*_2,33_ = 1.28, *P*=0.29; Genotype *F*_1,33_ = 0.001, *P*=0.96; 2-way ANOVA. **F)** GCs’ CNO induced normalized rate differences were plotted against their remapping index in Di^F/F^ controls and Ascl^CreER^;Di^F/F^ mice within a sliding 25 percentile-wide window of the normalized rate differences. GCs that are disinhibited by CNO have significantly lower remapping index in Ascl^CreER^;Di^F/F^ mice compared to those in Di^F/F^ controls. Pearson’s *R*_Cre-_=0.0241, *R*_Cre+_=-0.2744, \*\*\**P = 0*.*000928*, Fisher Z test. Error bars and shaded area, ± sem. *(See also Supplementary Figure 3)*

### Immature granule cells control rapid remapping of cue cells

We have recently described a functional heterogeneity in mGCs in response to sensory cue and self-motion information during spatial navigation (Tuncdemir et al., 2022). By controlling the administration of sensory cues and their association with locations on the treadmill track, we were able to isolate a population of ‘cue cells’, in addition to the classic population of ‘place cells’ recorded in the same sessions. Cue responses in single neurons were stable for long periods of time while place cells were more context selective (Tuncdemir et al., 2022). In the present study, we utilized similar cue manipulations to investigate whether iGCs contribute to this functional diversity within the granule cell population. In this task, the olfactory cue was omitted or shifted one-quarter of the track length (50 cm) once every 3-5 trials interleaved throughout the session (Figure 4A). Like in controls (data previously reported in (Tuncdemir et al., 2022)), in X-IR mice sensory cue responsive mGCs remapped their firing position to match the new location of the odor in cue-shift laps (Figure 4B, middle) and exhibited reduced activity in cue-omitted laps (Figure 4B, right), while the spatial representation of the track by place encoding mGCs were not affected by the odor cue. In both controls and X-IR mice, cue cells showed higher averaged spatial information, had more consistent spatial firing between the first and the second half of each session, and displayed earlier in-field onset firing than place cells (Figures S4A-C). However, compared to controls, X-IR mice displayed increased Ca^2+^ event rates both at the more frequent middle odor cue location and when the odor cue was presented at the infrequent “shift” location (Figures 4SD,E). Relatedly, while in control mice mGCs displayed smaller responses when the cue cells remapped upon shifting the cue to another location (Figure 4C); in X-IR mice the cue responses during the shift laps were comparable to the more frequent middle cue laps (Figure 4D). This resulted in enhanced remapping and reduced rate modulation of cue cells in X-IR mice compared to controls (Figure 4E). In agreement with the increased remapping of cue responses in X-IR mice, population correlations of firing rates of mGCs on middle cue and cue shifted laps was elevated at the odor cue location compared to controls (Figure 4F). In addition, these changes in mGC activity were limited to the cue responses as the Ca^2+^ event rates or population responses to omitting the cue were comparable between controls and X-IR mice (Figures 4G, 4SE). Thus, the absence of iGCs increases the remapping of cue cells and consequently reduces the influence of spatial location on cue responses. Taken together our results suggest that iGCs facilitate differential encoding of sensory cues based upon their spatial location by modulating the rate remapping of the cue cells when the cue is presented at different locations.

**Figure 4:**
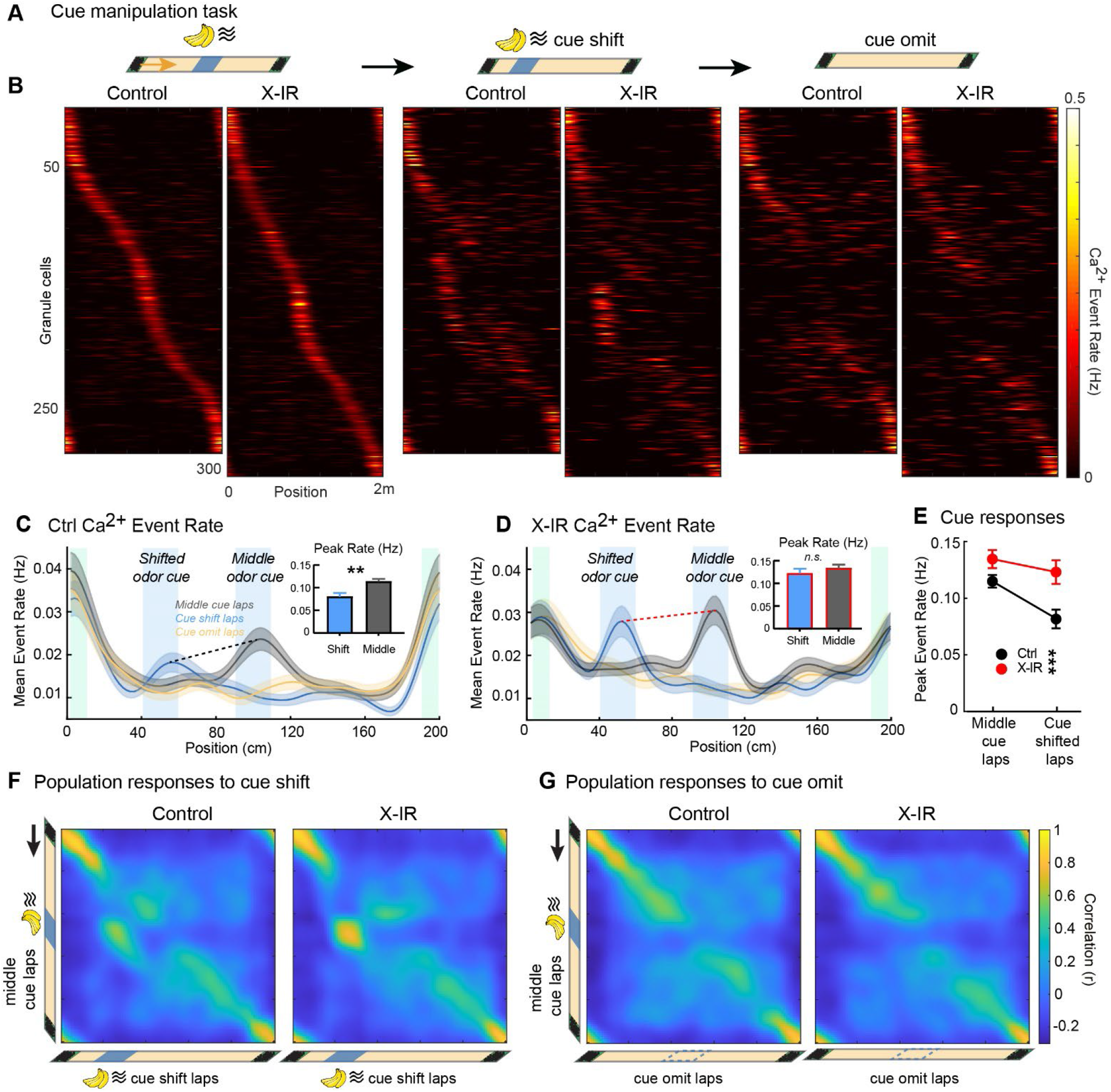
Chronic ablation of neurogenesis increases remapping of sensory cue responses. **A)** GC activity during the spatial cue task in control and X-IR mice. Location of the lap cue (green boxes) and an odor cue (blue box), on normal, and odor cue shift or omit laps. The two ends of an otherwise featureless treadmill belt taped on the opposite side of the RFID chip which served as the lap boundary cue. This cue was consistent through the testing of a mouse. Olfactory stimuli, an undiluted isoamyl acetate added to syringe filters, was delivered by opening a solenoid valve connected to a flow controller delivering constant airflow of compressed medical grade air for 1 sec. **B)** Lap-averaged spatial firing rates of spatially tuned neurons from controls and X-IR mice (n=8 mice) during the first session with the odor cue on normal middle location (left), cue-shifted (middle) and cue-omitted (right) laps. Each row represents activity of a single cell across lap types, sorted by activity on normal middle cue laps. N_Ctrl_=285 cells, 8 mice; N_IR_= 327 cells, 6 mice. **C)** Average spatial firing rates by position for neurons in control mice shown in panel “B” on normal (gray), cue-shifted (blue) and cue-omitted laps (yellow). (Inset) Averaged peak firing rate (Hz) of cue cells on laps in which the cue is shifted to the 50cm location and during normal middle (100cm) cue laps. *W* = -6371, \*\**P*_*Ctrl* – *IR*_ = 0.002, Wilcoxon rank sum test. **D)** Average spatial firing rates by position for neurons in X-IR mice shown in panel “B” on normal (gray), cue-shifted (blue) and cue-omitted laps (yellow). (Inset). Averaged peak firing rate (Hz) of cue cells on laps in which the cue is shifted to the 50cm location and during normal middle (100cm) cue laps *W* = -6044, *P*_*Ctrl* – *IR*_ = 0.0534, Wilcoxon rank sum test. **E)** Quantification of cue responses across genotypes. Peak firing of the cells responding to shifting of the odor cue is higher in X-IR mice compared to controls while the middle odor cue responses are similar in controls and X-IR mice. Interaction *F*_1,312_ = 0.18, *P*=0.18; Lap type *F*_1,312_ = 7.307, *P*=0.0072; Genotype *F*_1,312_ = 13.6, \*\*\**P*=0.0003; 2-way ANOVA. Planned comparisons: Middle cue laps: *P*_*Ctrl* – *IR*_ = 0.1177; Cue shift laps: \*\**P*_*Ctrl* – *IR*_ =0.0026; Tukey’s post hoc test. Control: N_OdorCueCells_= 78, X-IR N_OdorCueCells_= 80, Error bars, ± sem. **F)** Population vector (PV) correlations of granule cells with significant field at each treadmill position for cue shifted laps and normal middle cue laps in control (left) and X-IR (right) mice. For calculating PVs, lap-averaged spatial firing rate maps of all tuned cells in cue-shifted laps were correlated with that of normal middle cue laps. **G)** Population vector (PV) correlations of granule cells at each treadmill position for cue omitted laps and normal middle cue laps in control (left) and X-IR (right) mice (*see also Supplementary Figure 4)*

### Immature granule cells preferentially promote remapping of place cells during context discrimination

Having discovered that iGCs differentially influence place and cue cells, we revisited our initial experiments with different contexts (Figures 1 and 2) where multiple sensory features are used to define these contexts. We divided the spatially tuned population into place and cue encoding mGCs for subsequent analyses based on the position of their spatial fields during the context discrimination task. Place cells were defined as mGCs with peak Ca^2+^ event rates outside of the affixed tactile cues on the treadmill belt during the second exposure to context A (Figures 5A, S5A-B), while cue cells were defined as mGCs with peak activity within the boundary of the tactile cues on the belt (Figure 5B, S5A-B). Notably, there was a selective reduction in place rather than cue cell remapping in both models of iGCs inhibition. The high firing rate place cells in X-IR mice (Figure 5C) and the most disinhibited place cells in *Ascl*^*CreER*^ *Di* ^F/F^ mice (Figure 5E) showed the most pronounced deficit in remapping index. Conversely, following either chronic ablation (Figure 5D) or acute inhibition (Figure 5F) cue cells did not show significant differences in their remapping index compared to controls. Our results also demonstrate that the effects of CNO-induced inhibition of iGCs were most pronounced on the remapping of place cells that were disinhibited following iGC silencing (Figure 5G). Thus, our analysis suggests that iGCs selectively inhibit a subpopulation of place cells in *Di* ^F/F^ control mice that exhibit high context selectivity (Figure 5H). This relation between remapping and activity is not observed in cue cells (Figure S5C) or place cells that are not inhibited by iGCs during context discrimination (Figure S5D-E). Hence, our results suggest that iGCs enhance context selectivity by exerting a unique and differential effect on the remapping dynamics of place and cue encoding mGCs.

**Figure 5:**
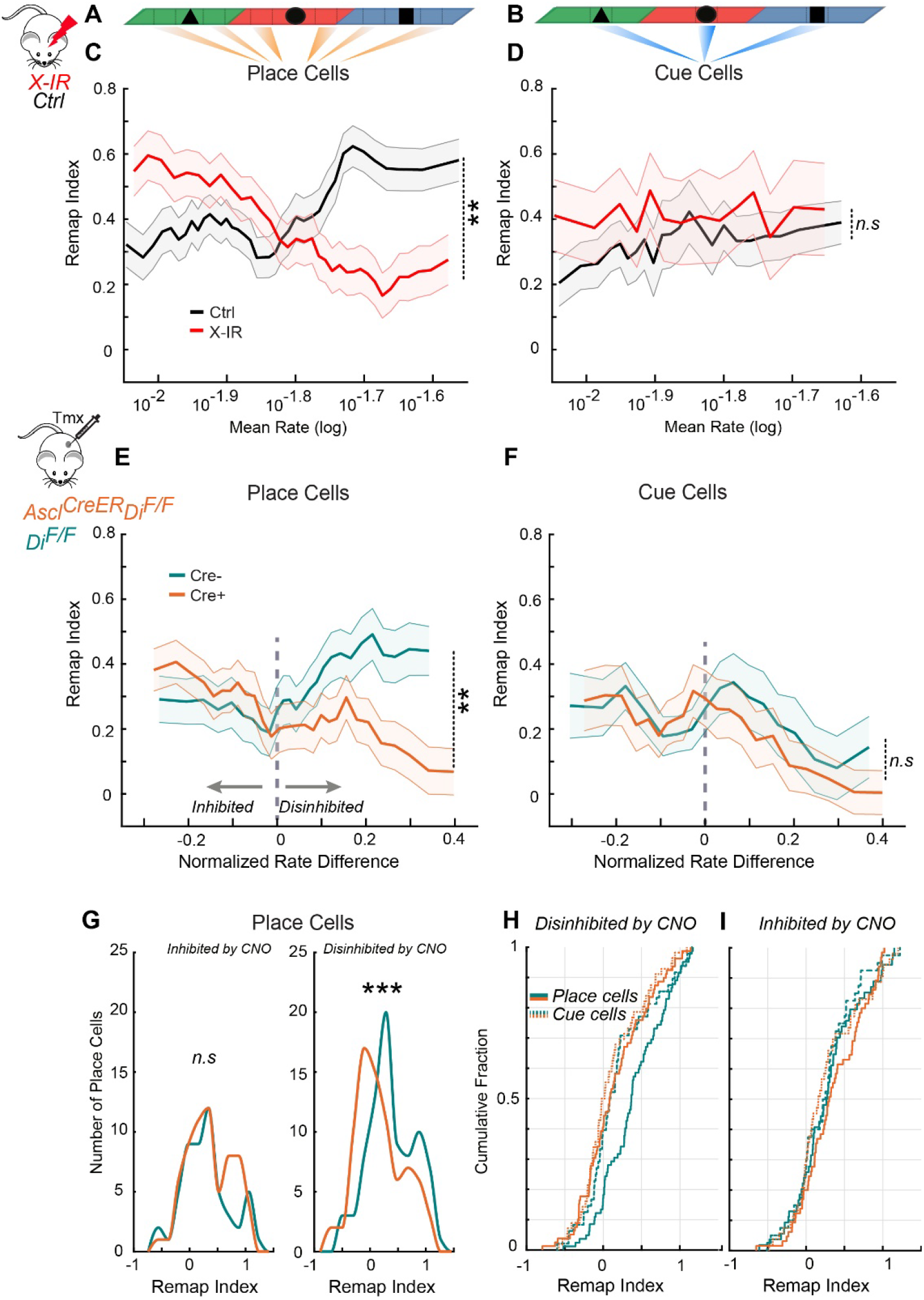
Immature granule cells preferentially promote global remapping of place cells during context discrimination. **A)** Significantly tuned population of mGCs during context discrimination tasks described in Figures 1,2,3 were divided according to peak Ca^2+^ event rate. Place encoding mGCs were defined as those with peak Ca^2+^ event rate > 5 cm from center of the affixed tactile cues on the treadmill belt during the second exposure to context A. **B)** Cue encoding mGCs were defined as those with peak Ca^2+^ event rate within the boundaries of the affixed tactile cues (10 cm wide) on the treadmill belt during the second exposure to context A. **C)** Averaged Ca^2+^ rates of place cells plotted against their remapping index in control (black) and X-IR (red) mice within a sliding window of the 25^th^ percentile of the averaged rates on a log scale. Context selectivity was calculated by subtracting the mean of the AB and A’B correlations from the AA’ correlation GCs with higher mean rates have significantly lower remapping index in X-IR mice compared to controls. Pearson’s *R*_Ctrl_=0.1561, *R*_IR_=-0.1754, \*\**P= 0*.*0012088*, Fisher Z test. **D)** Averaged Ca^2+^ rates of cue cells plotted against their remapping index in control and X-IR mice. Pearson’s *R*_Ctrl_=0.1560, *R*_IR_=0.0486, *P= 0*.*17888*, Fisher Z test. **E)** Place encoding mGCs’ CNO induced normalized rate differences were plotted against their remapping index in Di^F/F^ controls (green) and Ascl^CreER^;Di^F/F^ (orange) mice within a sliding window of the 25^th^ percentile of the normalized rate difference. Place cells that are disinhibited by CNO have significantly lower remapping index in Ascl^CreER^;Di^F/F^ mice compared to those in Di^F/F^ controls. Pearson’s *R*_Cre-_=0.0923, *R*_Cre+_=-0.2873, \*\**P= 0*.*0013236*, Fisher Z test. **F)** Cue encoding mGCs’ CNO induced normalized rate differences were plotted against their remapping index in Di^F/F^ controls and Ascl^CreER^;Di^F/F^ mice. Pearson’s *R*_Cre-_=-0.0321, *R*_Cre+_=-0.2582, *P= 0*.*11027*, Fisher Z test. Shaded areas, ± sem. **G)** The distribution of the context selectivity of place cells that are inhibited (left, *P*=0.48608) or disinhibited (right, \*\*\**P* = 0.00081) by CNO in Ascl^CreER^;Di^F/F^ mice compared to those in Di^F/F^ controls. Two-sample Kolmogorov-Smirnov test. **H)** Empirical cumulative probability distribution of the remapping index in place (straight lines) and cue (dotted lines) cells found in Ascl^CreER^;Di^F/F^ and Di^F/F^ mice based on their normalized rate responses to CNO. Note the excess fraction of disinhibited place cells in Di^F/F^ controls, which is lost on Ascl^CreER^;Di^F/F^ mice. *(See also Supplementary Figure 5)*

### Immature granule cells control long-term stability of DG representations

Lastly, we sought to characterize the influence of neurogenesis on the long-term stability of the GC subpopulations. We used the spatial cue task described above (Figure 4) to examine the responses of individual GCs on subsequent sessions with 1 hour between sessions or 1 week later (Figure 6A). We first used Bayesian decoding to infer how well we could predict the mouse’s location from firing rate vectors of mGCs that were cross-registered between sessions on the same day or 1 week later (Figure 6B*i*), regardless of their spatial tuning specificity. While the predicted position error was similar in both groups of mice when position was predicted using the cells found on sessions from the same day, the X-IR mice showed consistently lower prediction errors compared to controls when their position is predicted from neural data from 1 week later (Figure 6B*ii*). This resulted in a higher stability index in X-IR mice compared to controls (Figure 6B*iii*).

**Figure 6:**
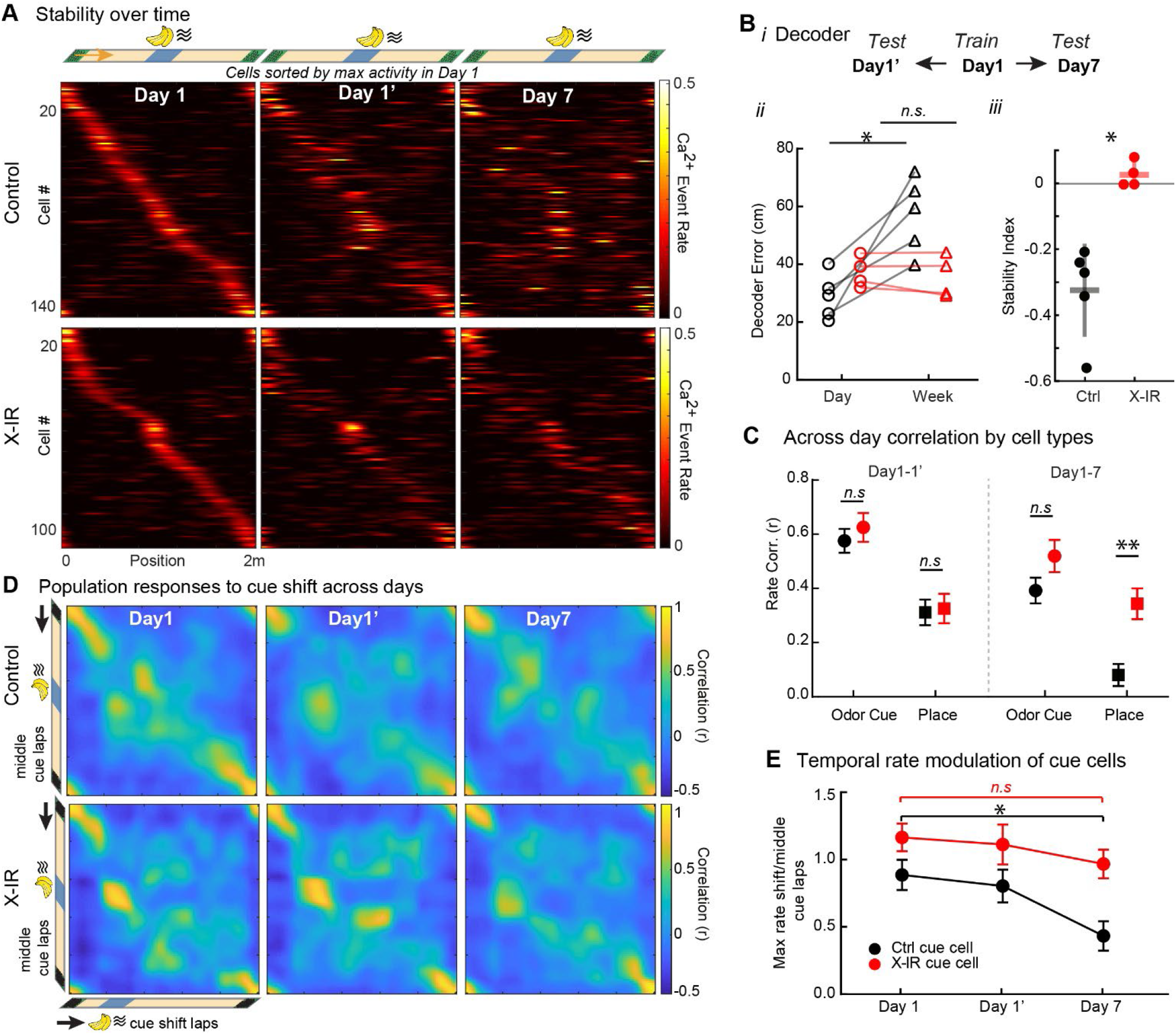
Chronic ablation of neurogenesis increases long-term stability of place cells and decreases rate modulation of cue cells. **A)** Spatial firing patterns of individual granule cells during the spatial cue task, matched between sessions within a day and over one week in control and X-IR mice. Top: Location of the odor (blue box) and lap cues (green boxes) on normal middle location laps. Bottom: Spatial firing rates for spatially tuned neurons tracked during subsequent sessions on the same day or one week later, ordered according to the position of peak activity during the first exposure on Day1. N_Ctrl_=144 cells, 5 mice; N_IR_= 106 cells, 4 mice. **B)** *i*-Position decoded by a Bayesian algorithm among cells found in all sessions (N_Ctrl_= 84.4 ± 10.5; N_IR_= 78.2 ± 7.2) by training using neural activity in the first session of Day 1 and calculating predicted position on the same day (Day1’) or one week later (Day7). ii-Line graphs represent median spatial decoder error calculated as the difference between the predicted position in the training data from those in test datasets that is higher in controls compared to X-IR mice when spatial activity from different days was used to decode position. *W* = -15, \**P*_*Ctrl-Day* – *Ctrl-Week*_ = 0.0313, *W* = -4, *P*_*IR-Day* – *IR-Week*_ = 0.625, Wilcoxon matched pairs signed rank test. iii. Stability index was calculated as: (Error _D1_-Error _D7_)/ (Error _D1_ + Error _D7_), *U* = 0, \**P*_*Ctrl-IR*_ = 0.0159, Mann-Whitney test. **C)** Mean spatial firing rate correlations within the same day and across 1 week for odor-cue cells and place cells in controls and X-IR mice. Interaction *F*_3,344_ = 2.73, \**P* = 0.043; Cell type *F*_3,344_ = 19.37, \*\*\**P*<0.001; Genotype *F*_1,344_ = 8.82, \*\**P* = 0.0032; 2-way ANOVA. Day1-1’: *P*^*Ctrl-Cue – IR-Cue*^ = 0.99; *P*^*Ctrl-Place – IR-Place*^ > 0.99; Day1-7: *P*^*Ctrl-Cue – IR-Cue*^ = 0.86; \*\**P*^*Ctrl-Place – IR-Place*^ = 0.0016; \*\*\**P*_*Ctrl-Cue* – *Ctrl-Place*_ = 0.0002; *P*_*IR-Cue* – *IR-Place*_ = 0.41; Tukey’s post hoc test. N_*Ctrl-Cue*_=39, N_*Ctrl-Place*_=65 from 5 mice, N_*IR-Cue*_=29, N_*IR-Place*_=47 cells from 4 mice. **D)** Gradual changes in population responses to cue shifts. Population vector (PV) correlations of lap-averaged spatial firing rate maps of cells with a significant field on Day 1 and found on the same day (Day1’) or one week later (Day7) in cue-shifted laps were correlated with that in normal middle cue laps, calculated at each treadmill position in control (top) and X-IR (bottom) mice. **E)** Temporal rate modulation of cue cells, calculated as the ratio of peak rate during cue shifted laps relative to the more frequent middle cue laps for all cue cells on days 1, 1’ and 7. Note that lower values in cue cells found in Control mice (black) mean smaller responses cue shifting, while cue responses remain elevated in X-IR mice (red). Interaction *F*_2,186_ = 0.65, *P* = 0.52; Day *F*_2,186_ = 3.95, \**P*=0.021; Genotype *F*_1,186_ = 14.06, \*\**P* = 0.0002; 2-way ANOVA. Planned comparisons: Ctrl: \**P*_*Day1* – *Day7*_ = 0.032; IR: *P*_*Day1* – *Day7*_ = 0.93, Tukey’s post hoc test. N_*Ctrl-Cue*_=39, N_*IR-Cue*_=29 cells. Error bars, ± sem.

We then examined cue and place populations separately to determine the relative influence of iGCs on these two populations. In control mice, cross-registered odor cue cells fired reliably to the same olfactory cue after 1 week, while the correlations in the activity of place cells after 1 week were significantly lower (Figure 6C, (Tuncdemir et al., 2022)). In X-IR mice however, place cells displayed reduced remapping from Day 1 to Day7 compared to controls, as measured by increased correlation of their tuning curves (Figure 6C). Further, when we examined how cue representations change over time, we observed an increase in remapping to the cue shifts 1 week later in X-IR mice compared to controls, as measured by population correlations of firing rates of mGCs on middle cue versus shifted cue laps (Figure 6D). This is consistent with higher levels of cue cell activity in the cue shifted laps relative to the middle cue laps after 1 week in X-IR mice (Figure 6E). Thus, both across time and in different contexts, iGCs bidirectionally modulate remapping of cue and place cells, whereby they limit rate modulation of cue cells but promote the flexibility of place cells, which results in improved pattern separation.

## Discussion

Using *in vivo* two-photon Ca^2+^ imaging, we assessed changes in mGC spatial and contextual representations following either chronic ablation or acute inhibition of iGCs. A key property of mGCs is their ability to change their firing fields in response to changes in the surrounding environment, a phenomenon called remapping (Colgin et al., 2008). Here, we report that the activity of synaptically immature but functionally integrated adult born iGCs directly control the remapping of mGCs firing fields in response to environmental or contextual changes thereby decreasing the overlap of the active ensembles representing each context. While the pattern separation function of the iGCs and their ability to promote the formation of non-overlapping ensembles representing each context has been alluded to previously (McAvoy et al., 2015; Johnston et al., 2016; Tuncdemir et al., 2019), our study provides the first *in vivo* physiological evidence of the role of the iGCs in pattern separation. Further, the aberrantly increased stability of the contextual representations resulting from ablation or inhibition of iGCs is not due to a broad disinhibition of mGCs or changes in their spatial tuning. Our data suggest instead that the modulatory impact of iGCs depends on the specific contextual features the mGCs encode, such as information about the sensory cues within an environment versus mice’s self-motion information in between these cues. Specifically, our findings support a model where the activity of the iGCs improves pattern separation by promoting remapping of place cells (Figure S6A) and limiting remapping of cue cells (Figure S6B). This likely contributes to pattern separation by allowing the formation of distinct traces when the same cues or objects are encountered in different locations. On the other hand, the absence of iGCs prevents the flexible representations of different contexts by overemphasizing sensory cue representations shared between contexts at the detriment of spatial features (Figure S6C). The ability of iGCs to bidirectionally control remapping depending on the type of information mGCs encode, likely amplify their influence and underlie how a small number of iGCs constituting less than 5% of all GCs (Mathews et al., 2010) exert an outsized influence on DG circuit function and behavior.

### Adult born granule cells modulate mGCs

Examining the activity of genetically identified iGCs *in vivo* has revealed that they are highly excitable and display a lower spatial tuning specificity than mGCs, features which may be ill suited for coding spatial information (Danielson et al., 2016; McHugh et al., 2022). Here, we extend these observations and provide evidence in support of a modulatory rather than a direct role of iGCs in flexible encoding of different contextual features. Our findings reveal that chronic or acute manipulations of iGCs do not impair initial encoding of contexts A or B (Figures 1,S1,2,S2), and that population level mean firing rates as well as spatial information content are comparable across groups of mice (Figure 3, S3). In addition, we had shown earlier that iGCs and mGCs exhibit similar levels of remapping during sequential exposures to distinct treadmill environments (Danielson et al., 2016). Thus, the selective impairments in the remapping of mGCs we observe here after inhibiting iGCs is consistent with the loss of a modulatory rather than an instructive role for iGCs. Furthermore, the functional segregation of the mGC population into cue and place encoding populations is intact in the absence of iGCs (Figures 4,S4). Hence, while the details of the contextual features such as the sensory cue and self-motion information can emerge and be refined during cognitive map formation without iGCs, the ability of the DG network to flexibly form new representations during subsequent exposures to different contexts is impaired.

### Origins of iGC mediated remapping in the DG

Extensive prior work has proposed that the inputs from the entorhinal cortex trigger hippocampal remapping (Colgin et al., 2008; Latuske et al., 2017). The main long-range excitatory inputs of the mGCs come from the medial and lateral entorhinal cortices (MEC and LEC, respectively) (Witter, 2007). The MEC is thought to prominently encode self-motion information, while the LEC is thought to primarily represent information about sensory cues (Hargreaves et al., 2005; Knierim et al., 2014). Global remapping of CA1/3 place cells, whereby place cells change their preferred firing location, is strongly associated with the activity of the grid cells in the MEC (Fyhn et al., 2004; Miao et al., 2015; Rueckemann et al., 2016). Our finding that iGCs inhibit a subpopulation of place cells that exhibit global remapping (Figures 5,S5) suggests that this iGC-mediated inhibition is necessary for the emergence of new place fields possibly as a result of new MEC grid cells inputs (Figure S6A). Such inhibition may be triggered by local inhibitory interneurons (Ikrar et al., 2013; Temprana et al., 2015; Drew et al., 2016). On the other hand, remapping can occur even without inputs from the MEC (Schlesiger et al., 2018) and GCs also receive inputs from the supramammillary nucleus of the hypothalamus (Farrell et al., 2021; Li et al., 2022) and the lateral septum (Ogando et al., 2021; Gómez-Ocádiz et al., 2022). It is therefore possible that the MEC is not the only input responsible for the impact of iGCs on place cell remapping.

The circuit mediating the impact of iGCs on the remapping of cue cells likely involves the LEC. Notably, a recent study has demonstrated specific impairments in hippocampal rate remapping upon lesions of the LEC (Lu et al., 2013). Furthermore, iGCs receive strong monosynaptic inputs from the LEC during their hyperplastic phase (Vivar et al., 2012; Woods et al., 2018), which in turn reduce the activity of mGCs in response to LEC inputs (Luna et al., 2019). Hence, we suggest that one function of iGCs is to modulate LEC inputs to cue cells either via interneurons (Ikrar et al., 2013; Temprana et al., 2015; Drew et al., 2016) or via direct monosynaptic inhibition of cue cells through group II metabotropic glutamate receptors (Luna et al., 2019). This inhibition may promote rate modulation of the cue cells induced by changes in their location (Figures 4,6) to promote the formation of distinct ‘rate modulated’ traces for the same cues encountered in different locations (Figure S6B).

### Neurogenesis and overgeneralization

The phenotype displayed by the X-IR and *Ascl*^*CreER*^ *Di* ^F/F^ mice may be characterized by an overreliance on specific cues at the detriment of spatial locations when encoding novel environments (Figure S6C). This phenotype is reminiscent of the overgeneralization that is often observed in anxiety disorders such as PTSD (Lissek et al., 2010; Kheirbek et al., 2012; Lange et al., 2017). For example, a patient who developed PTSD following 9/11 may display an exaggerated fear response at the sight of a plane or a tower because of an inability to separate the present safe situation from the past traumatic experience. It will therefore be interesting to investigate whether strategies aimed at modulating DG function either indirectly by stimulating neurogenesis or directly by modulating the activity of mGCs may be beneficial for the treatment of patients who display this form of overgeneralization.

## Acknowledgements

This work was funded by NIH K99 MH12226 to S.N.T., Revson Fellowship in Biomedical Science to A.D.G., K01 AG054765 to V.M.L.; R37 MH068542, R01 MH083862, R01 NS081203 to R.H., the Hope for Depression Research Foundation (HDRF RGA-13-003) to R.H., NYSTEM (C029157) to R.H., NIMH R01 MH100631, NINDS R01NS094668, and NINDS U19NS104590 (A.L.). We would like to thank S. Dymecki for providing us with the Cre-responsive *loxP-stop-mCherry-loxP-hM*_*4*_*D*_*i*_ mouse line.

## Author Contributions

S.N.T. and R.H. designed the research and wrote the paper. S.N.T. performed experiments and analyzed the data with assistance from A.D.G., H.C and C.O.L. V.M.L. conducted in vitro electrophysiological experiments. A.L., A.D.G., V.M.L. and C.O.L provided technical infrastructure and input to the manuscript.

## Methods

### Resource Availability

#### Lead Contact

Further information and requests for resources and reagents should be directed to and will be fulfilled upon reasonable request by the lead contact.

##### Materials availability

This study did not generate new unique reagents.

#### Data and code availability

All data reported in this paper will be shared by the lead contacts upon request. Any additional information required to reanalyze the data reported in this paper is available from the lead contact upon request.

### Mice

All procedures were conducted in accordance with the U.S. NIH Guide for the Care and Use of Laboratory Animals and the Institutional Animal Care and Use Committees of New York State Psychiatric Institute and Columbia University. Adult male and female C57BL/6J mice were supplied by Jackson Laboratory for irradiation experiments, *Ascl-creERT2*^+/−^ were purchased form Jackson Laboratory and bred in house with *loxP*-*stop-mCherry-loxP-hM4Di* mice to obtain male and female Cre^−^ and Cre^+^ littermates from heterozygote breedings. At least three separate cohorts of mice were run for all experiments. Mice were housed in a vivarium grouped 2-4 mice/cage enriched with running wheels, maintained on a 12-hour light cycle and used at 8-10 weeks of age. To activate Cre-recombinase activity, mice at 8-10 weeks of age were administered *i*.*p*. tamoxifen injections for 5 consecutive days (100 mg kg^−1^ suspended in corn oil). The 5th day was designated post-induction day 0. Imaging experiments were conducted 4-5 weeks later. For in vivo silencing of iGCs in *Ascl*^*CreER*^ *hM4Di* ^F/F^ compared to *Cre* negative *hM4Di* ^F/F^ littermates CNO was dissolved in 100% dimethyl sulfoxide (DMSO) and diluted with 0.9% saline to a final concentration of 5 μM CNO and 0.001% DMSO. 30-60 minutes before the imaging session all mice received 5 mg kg^−1^ CNO diluted in 0.9% saline, administered by *i*.*p*. Experiments were conducted during the light portion of the cycle. Food and water were available *ad libitum* until the beginning of the experiment, when they were placed under controlled water supply and maintained at >90% of their pre-deprivation weight over the course of imaging experiments.

### Focal X-irradiation

Hippocampal irradiation was conducted as previously described (Burghardt et al., 2012; Denny et al., 2012). Briefly, after ketamine (105 mg/kg, i.p.) and xylazine (7 mg/kg, i.p.) anesthesia, 8–10-week-old mice were placed in a stereotaxic frame, and cranial irradiation applied. There was a 3.22 × 11-mm window above the hippocampus in a lead plate that otherwise shielded the entire body and allowed for focal Xray application. X-rays were filtered using a 2 mm Al filter, the corrected dose rate was approximately 1.8 Gy per min and the source to skin distance was 30 cm. A cumulative dose of 5 Gy was given over the course of 2 minutes and 47 seconds on 3 days with 2-3 days between each X-ray session. Control mice did not receive radiation, but were housed, anesthetized, and transported with irradiated mice throughout the experiment. Injection and implantation were performed 8 weeks later.

### Surgery

Dentate gyrus virus injection and imaging window implantation surgeries were performed as described previously (Tuncdemir et al., 2022). For all surgical procedures, mice were anesthetized with 1.5% isoflurane at an oxygen flow rate of 1 L/min, and head-fixed in a stereotactic frame (Kopf Instruments, Tujunga, CA). Eyes were lubricated with an ophthalmic ointment, and body temperature maintained at 37°C with a warm water recirculator (Stryker, Kalamazoo, MI). The fur was shaved and incision site sterilized prior to beginning surgical procedures, and subcutaneous saline and carpofen were provided peri-operatively and for 3 days post-operatively to prevent dehydration and for analgesia. Mice were unilaterally injected with recombinant adeno-associated virus (rAVV) carrying the GCaMP6s or GCaMP7s transgenes (pAAV.Syn.GCaMP6s.WPRE.SV40 or pGP-AAV-syn-jGCaMP7s-WPRE) purchased from Addgene (viral preps #100843-AAV1, 104487-AAV1) with titers of 1-5×10^12^ in dorsal dentate gyrus using a Nanoject syringe (Drummond Scientific, Broomall, PA). Unilateral injection coordinates for dorsal DG were -2 mm AP, -1.5 mm ML, and -1.85, -1.7, -1.55 mm DV relative to the cortical surface 30 nL of 1:3 diluted virus was injected at each DV location in 10 nL increments. Mice were allowed to recover for 3 days and then were unilaterally implanted with an imaging window and stainless-steel head-post for head fixation. Imaging windows were constructed by adhering 2 mm diameter, 2.3 mm long stainless-steel hypodermic tubing (Ziggy’s Tubes and Wires Inc, Pleasant Hill, TN) to 2 mm diameter glass coverslips (Potomac Photonics, Halethorpe, MD). A 2 mm diameter craniotomy was made centered on the previous injection site with a taper pointed-drill (Henry Schein Inc, 9004367) and dura was removed with micro curette (FST, 10080-05). The overlying cortex was gently aspirated to reveal capsular fibers with continuous irrigation with ice cold aCSF solution and bleeding was controlled with a collagen gel sponge (Avitene). Under minimal bleeding, a 30g blunt syringe was used to gently aspirate capsular and CA1 alveus fibers with white appearance and CA1 pyramidale and moleculare with pink appearance until vasculature of the hippocampal fissure became visible (under bright light with low bleeding). The cannula, attached to the stereotactic handle, was then gently lowered into the craniotomy and affixed to the skull using dental cement (Unifast Trad powder and LC light cured acrylic UV, Henry Schein).

### In vitro electrophysiology

Chemogenetic manipulation of iGCs using *Ascl*^*CreER*^ *hM4Di* ^F/F^ mice was confirmed by using *in vitro* whole-cell current clamp recordings on mGCs in slice preparations as described previously (Luna et al., 2019). Briefly, after 4-6 weeks post-tamoxifen inductions, mice were anesthetized by isoflurane inhalation, decapitated, and brains rapidly removed. DG slices (350μm) were cut on a vibratome (Leica VT1000S) in ice cold partial sucrose artificial cerebrospinal fluid (ACSF) solution (in mM): 80 NaCl, 3.5 KCl, 4.5 MgSO4, 0.5 CaCl2, 1.25 H2PO4, 25 NaHCO3, 10 glucose, and 90 sucrose equilibrated with 95% O2 / 5% CO2 and stored in the same solution at 37°C for 30 minutes, then at room temperature until use. Recordings were made at 30-32°C (TC324-B; Warner Instrument Corp) in normal ACSF (in mM: 124 NaCl, 2.5 KCl, 1 NaH2PO4, 25 NaHCO3, 20 glucose, 1 MgCl2, 2 CaCl2). Whole-cell recordings (−70 mV) were obtained using a patch pipette (4.5-6.5 M) containing (in mM): 135 KMethaneSulfate, 5 KCl, 0.1 Na-EGTA, 10 HEPES, 2 NaCl, 5 ATP, 0.4 GTP, 10 phosphocreatine (pH 7.2; 280-290 mOsm). Recordings were made without correction for junction potentials. Granule cells were visualized and targeted via infrared-differential interference contrast (IR-DIC; 40x objective) optics on an Axioskop-2 FS (Zeiss). For perforant path stimulation, a concentric bipolar stimulating electrode (FHC) controlled by a stimulus isolator (ISO-flex, AMPI Instruments) was positioned on the DG molecular layer (triggered at 0.04 Hz). Perforant path inputs were stimulated at 40 Hz (20 pulses every 25 ms) at the lowest stimulation intensity to elicit 100% responses in mGCs. Current and voltage signals were recorded with a MultiClamp 700B amplifier (Molecular Devices, USA), digitized at 5–10 kHz, and filtered at 2.5–4 kHz. Data were acquired and analyzed using Axograph (Axograph Scientific, Sydney, Australia). Evoked synaptic responses were quantified by calculating the AUC (cumulative area above (+) and below (−) the baseline in mV s).

### Behavioral training and apparatus for head-fixed imaging

After a minimum of 1 week recovery period, mice underwent a water restriction scheme (1ml per day) and trained to run on treadmill while head-restrained. The training period typically lasted a week (2 training sessions/day, 15 min each) until the mice were able to run for at least 1 lap/ minute and seek reward from one of 3 reward zones that were randomly selected along the belt on each traversal by licking the water delivery port. During the training period mice ran on a cue-less ‘burlap’ belt and progressed to a different belt containing cues of different modalities as described below. For context discrimination experiments mice ran at will and for cue manipulation experiments separate cohorts of mice ran on a motorized belt as described previously (Tuncdemir et al., 2022). For mice running on motorized treadmill, we initiated the motorized belt after mice were able to run for at least 1 lap/ minute, adjusted to the natural velocity of each mouse and proceeded training for 1-2 more days. We discarded mice that did not perform sufficiently to receive getting all of their daily water supply during treadmill training, and were not motivated to move on the treadmill. Mice were imaged for three or two consecutive sessions/day, each 15 minutes long, for the duration of 7-10 days. Contextual representations described in Figures 1-3, 6, S1-4 5 and single odor responses described in Figures 4, 5 and S5 are from the first-time mice were exposed to these contexts or sensory stimuli, respectively. Additionally, in all experiments, the treadmill belt material was changed between sessions to reduce the chances of urine contamination which might act as an additional olfactory cue. In between experiments, belts were washed with Tergazyme (Alconox).

The behavioral apparatus consisted of 2m long, 3” wide cotton fabric belt stretched between 6” diam. laser-cut plastic wheels, mounted on an aluminum frame (8020.net). Spatial triggering of task events was performed by custom software via serial communication with a microcontroller (Arduino DUE) and an associated printed circuit board (OpenMaze OM4 PCB, www.openmaze.org) on the treadmill. The axle of the treadmill wheel was attached to a quadrature rotary encoder (US Digital #MA3-A10-125-B) connected to a custom quadrature decoder board and Arduino Nano (courtesy of Wen Li). Angular displacement was converted into a virtual linear distance based on the circumference of the treadmill wheels, and laps were determined by reading an RFID chip on the treadmill belt attachment point with an RFID reader (Sparkfun #ID12LA) mounted under the animal’s head fixation point. A water reservoir connected to a water delivery port consisting of a small gavage needle (Cadence Science) was placed within reach of the mouse’s tongue. A capacitance touch sensor (Sparkfun MPR121) was attached to the water port to measure licking and the sensor was connected to the Arduino/OM4 PCB. Small 2-3ml drops of water were delivered by the brief opening a solenoid valve (Parker Hannefin) connected to the water port. Rewards were triggered at random locations each lap when mice entered a 10cm long reward zone on the track and were available until mice exited the reward zone or 3 sec had elapsed. For contexts, similar to our previous work, each context (A and B) consisted of the same treadmill belt (the same sequence of three joined fabric ribbons) but was distinct in its visual, auditory, tactile, and olfactory stimuli (Figure S5A,B, (Danielson et al., 2016)). To allow for comparison of GC activity between similar contexts, the same three fabrics were used in the same order, but the locations of all of the tactile cues were shuffled between the two belts. For X-irradiation experiments olfactory cue was delivered ambiently inside the rig similar to our previous work (Danielson, 2016), while for DREAAD experiments, undiluted isoamyl acetate (banana odor, Sigma W205532) or menthalactone (mint odor, Sigma W376418) were added to syringe filters (Whatman #6888-2527) and delivered by opening a solenoid valve (SMC) connected to a flow controller delivering constant airflow of compressed medical grade air for 1s (∼3psi). For cue manipulation task, undiluted isoamyl acetate (banana odor) was also delivered in 1s pulses at indicated locations. Custom written Be-Mate algorithm implemented in Java was used for recording mice’s licking, its position on the belt, and cue delivery. Mice were monitored using an IR camera (PS3eye) and illuminated using an IR LED array. For motorized running, a 12V DC gear motor was attached to the axle of the treadmill connected to a separate Arduino/OpenMaze shield using pulse-width modulation to adjust the rotation speed.

To isolate cue-selective responses among the granule cell population, normal cue laps in which the olfactory, cue was presented in the middle of the treadmill track (90-110cm) were interspersed with occasional laps (10-20% of laps) in which the same cue was omitted (“omit” laps), or shifted forward ¼ of the track (“shift” laps, 40-60cm cue position).

### In vivo two-photon imaging

Imaging was conducted using a microscope setup which consists of 8kHz resonant galvanometer (*Bruker*) mounted to a mirror-based multi-photon microscopy system (*Prairie Technologies*) and an ultra-fast pulsed laser beam (920-nm wavelength; *Chameleon Ultra II, Coherent*, 20–40-mW average power at the back focal plane of the objective) controlled with an electro-optical modulator (*Conoptics*, Model 302 RM). GCaMP fluorescence was excited through a 40x water immersion objective (Nikon NIR Apo, 0.8 NA, 3.5 mm WD) and fluorescence signals detected with photomultiplier tubes (*Hamamatsu 7422P-40*), acquired with PrairieView software (*Prairie*) at 30fps frame rate (512×512 pixels, 1.3 mm/pixel). A custom dual stage preamp (1.4×10^5^ dB, Bruker) was used to amplify signals prior to digitization. Two goniometers (Edmund Optics) were used to adjust the angle of each mouse’s head in order to achieve the same imaging plane over multiple sessions.

#### Pre-processing for Ca^2+^ imaging

Movies were motion corrected using NoRMCorre algorithm using a non-rigid registration method that splits the field of view (FOV) into overlapping patched that are registered separately then merged by smooth interpolation (Pnevmatikakis and Giovannucci, 2017). Videos were then spatially and temporally down-sampled by 2 to reduce noise and the computational power required for cell segmentation. Spatial and temporal components for individual cells were extracted using the singular value decomposition method by *Suite2p* algorithm (https://github.com/cortex-lab/Suite2P), followed by visual inspection of the regions of interest (ROIs) using the Suite2p graphical user interface to manually select small, densely packed DG granule cells and discard large isolated cell bodies corresponding to mossy cells or other hilar interneurons. To obtain total number of DG granule cells within the imaging fields of view in a subset of total sessions, time averaged images were segmented and counted using the *Cellpose* algorithm (https://github.com/MouseLand/cellpose) followed by manual inspection to discard large cell bodies corresponding to mossy cells or other hilar interneurons.

#### Transient detection

Ca^2+^ transients were detected and used for all subsequent analysis. Transient detection was done with a threshold of height based on an estimate of the noise as the standard deviation of the signal. Ca^2+^ transient events were defined as fluorescence peaks with a rise slope greater than 4 standard deviations above an iteratively refined baseline that last for at least 3s.

#### Behavioral and Calcium Data Alignment

Behavioral data was aligned to Ca^2+^ data using the record of a synchronization signal between the two computers used for data collection. Behavioral data was down-sampled to match Ca^2+^ imaging data.

### Data Analysis

Data were analyzed using custom-written routines implemented in MATLAB. Plots were generated in MATLAB and Prism.

#### Identification of spatially-tuned neurons

We restricted our analysis to continuous running at least 2 sec in duration and with a minimum peak speed of 5 cm/sec. For each lap crossing, position data and Ca^2+^ transient events for each cell were binned into 2 cm-wide windows (100 bins), generating raw vectors for occupancy-by-position and calcium transient numbers-by-position which were then circularly smoothed with a Gaussian kernel (*SD* = 5 cm). A firing rate-by-position vector was computed by dividing the smoothed transient number vector by the smoothed occupancy vector. Within each lap, we circularly shuffled the positions 1000 times and recomputed firing rate-by-position vectors to generate a null distribution for each spatial bin. A spatially selective cell was defined that met the following criteria: (a) the cell should fire above its mean firing rate within its spatial field in at least 20% of laps or for a minimum of 3 laps; and (b) observed firing should be above 99% of the shuffled distribution for at least 5 consecutive spatial bins (10 cm) wrapping around the two edges of the belt. For cue manipulation experiments, we have identified spatially tuned neurons by excluding bins in which sensory cues were omitted or shifted and calculated firing rate vectors in these laps separately. For context discrimination experiments, among all the spatially tuned cells, cue cells were defined as those with peak spatial field rate that was within 15^th^ and 20^th^ bins, 50^th^ and 55^th^ bins, 80^th^ and 85^th^ bins corresponding to the tactile cues (or the odor cue which was delivered within 50^th^ and 55^th^ bins for CNO experiments). The remaining cells constituted the “place cells”. For cue manipulation experiments, “*middle cue cells*” (also termed as odor-cue) were defined as those with peak amplitude that was within 50^th^ and 55^th^ bins and were at least two times larger than those in cue-omitted laps in the same session. These experiments also had “lap-cue cells” due to prominent texture of the edges of the belt and were defined as those with averaged spatial fields overlapping at least 50% of the region wrapping around the 90^th^ and 10^th^ bins in the normal laps that were treated separately. The remaining cells constituted the “place cells”.

#### Spatial information, stability, consistency, and emergence of spatial fields

To calculate a measure for spatial information content for granule cells we adapted a traditional method of spatial information assessment (Skaggs et al., 1993; Danielson et al., 2016; Grosmark et al., 2021) to Ca^2+^ imaging data. For each cell, we used the firing rate-by-position vector and shuffled null distribution computed above and calculated the spatial information content as follows:

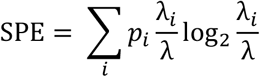

Where λ is the overall mean event rate, λ_*i*_ is the mean event rate of each cell in the *i*^th^ bin, *p*_*i*_ is the probability of that mouse being in the *i*^th^ bin that is occupancy in the *i*^th^ bin/ 15mins.

To account for the fact that low firing rates artificially produce high spatial information scores, we subtracted the mean of the shuffled information per spike from observed information per spike, divided by the standard deviation of the shuffled values to determine the spatial variance for each cell. Therefore, the amount of spatial information is inferred from differences in normalized Ca^2+^ activity in each neuron and reported as bits per seconds. The consistency of place field firing was determined as the cross-correlation between the averaged firing rate-by-position vector of the first and the second halves of the total number of cue normal laps within a session. We determined place field onset lap in cue normal laps as described previously (Sheffield et al., 2017). Briefly, starting on lap 1 we searched for a significant Ca^2+^ transient event present within the boundaries of the previously determined mean spatial field calculated from all the laps in the session. If one were found we would then search for Ca^2+^ transient event on each of the next 4 laps. If 3 of the 5 laps had Ca^2+^ transients within the mean place field boundaries, lap 1 would be considered the place field onset lap. If either lap 1 had no Ca^2+^ transient or less than 3 of the 5 laps had Ca^2+^ transient, we would move to lap 2 and repeat the search.

#### Multi-Session Cell Tracking

Cells were tracked across sessions using CellReg algorithm reported in (Sheintuch et al., 2017). Briefly, rigid alignment with both translations and rotations was performed on spatial footprint projections of each session and manually inspected for quality. To improve performance with our data, we modified the CellReg source code to consider complete spatial footprints instead of centroids during alignment. The centroid distance between neighbors was then calculated and used to create a probabilistic model that estimated the expected error rate at different thresholds. The optimal centroid distance threshold was chosen by the algorithm and used to match cells. A clustering algorithm then refined these decisions previously made using pairwise comparisons.

Following cell registration, tracked cells were matched with their corresponding functional cell types (i.e. Spatially tuned cell, non-spatially tuned cell for Figure S3B,C, E,F or mid-cue, place cells for Figure 6D, as described above). The analyses presented in Figures S3 and 6 are carried out pairwise, to maximize the number of cells in each comparison and to minimize the total number of comparisons. For context comparisons we used Context A’ and for multiday comparisons we used Day1 as the reference sessions. To calculate the fraction of cells that maintain their identity, cell pairs that were counted as being the same cell type in both sessions was divided by all of that cell type in the reference session. In order to derive a null distribution for preservation of pairwise identity, we randomly permutated the cell IDs of all the tracked cells in pairwise sessions 1000 times and calculated the fraction of cells that were the same, among all of that cell type in the normal session. We calculated p-values by comparing actual data to this null distribution, 97.5^th^% of the null distribution is presented dotted lines in Figure S3 B,C, E,F.

#### Rate correlation, population vector (PV) and remapping index analysis

Comparison of single cell activity between different sessions was calculated using Pearson’s correlation of the spatially binned, averaged firing rate-by-position vector in different context sessions in Figures 1,2 and across day sessions in Figure 6. The variability in neural population activity between different contexts (Figures S1, S2, 6) was calculated by using Pearson’s correlation on each 2 cm bins of the firing rate-by-position vector along the treadmill. In Figure 4 and 6, spatial firing rates for each spatially tuned cell on normal middle cue laps were cross-correlated with firing rates on shift laps on each 2 cm bins of the firing rate-by-position vector along the treadmill in order to visualize the cross-correlation peak offset (i.e. rate remapping) after cue manipulation. Remapping index in Figures 3, 5 and S5 were calculated by subtracting the mean averaged correlation between different contexts (Contexts AB and A’B) from the correlation between similar contexts (sequential exposures to Context A, Contexts AA’).

#### Bayesian Reconstruction Analysis

To calculate the probability of the animal’s position given a short time window of neural activity, we used a previously published method based on Bayesian reconstruction algorithm (Davidson, Wilson 2009, Grosmark and Buzsáki, 2016). Briefly, Ca^2+^ transient events for each cell were binned into 1 second windows to construct firing rate vectors. For each of these binned firing rate vectors, Bayesian classification of virtual position (posterior probability for each bin) was performed utilizing a template comprising of a cell’s smoothed firing rate-by-position vectors. In order to cross-validate our decoding procedure, we used the firing rate-by-position from a cell that were cross-registered from one session (for example Context A and B or Day 1’ and Day7) as “testing” dataset while the that cell’s firing rate-by-position vector in another session (for example Context A’ or Day1, respectively) constituted the “training” dataset. The resulting posterior probability distribution for each bin is the likelihood for an animal is located in that bin, which adds up to 1, and the bin with the maximum posterior probability is the estimated position of the animal. Post-reconstruction, we divided the time bins (excluding those with no activity) and according to the lap types when necessary (excluding omit or shift laps). To determine the decoding error, we calculated the absolute difference between the animal’s actual position and the maximum posterior probability in that bin. For all decoding of the decoding analyses, we used all of the cells cross-registered across sessions regardless of their tuning profiles. To determine the discrimination index, we subtracted the decoder error calculated by subtracting Context A from that of Context B and divided by the sum of these values. To determine the stability index, we subtracted the decoder error of Day7 from that of Day1’ and divided by the sum of these values.

## Figures

**Supplementary Figure 1.**
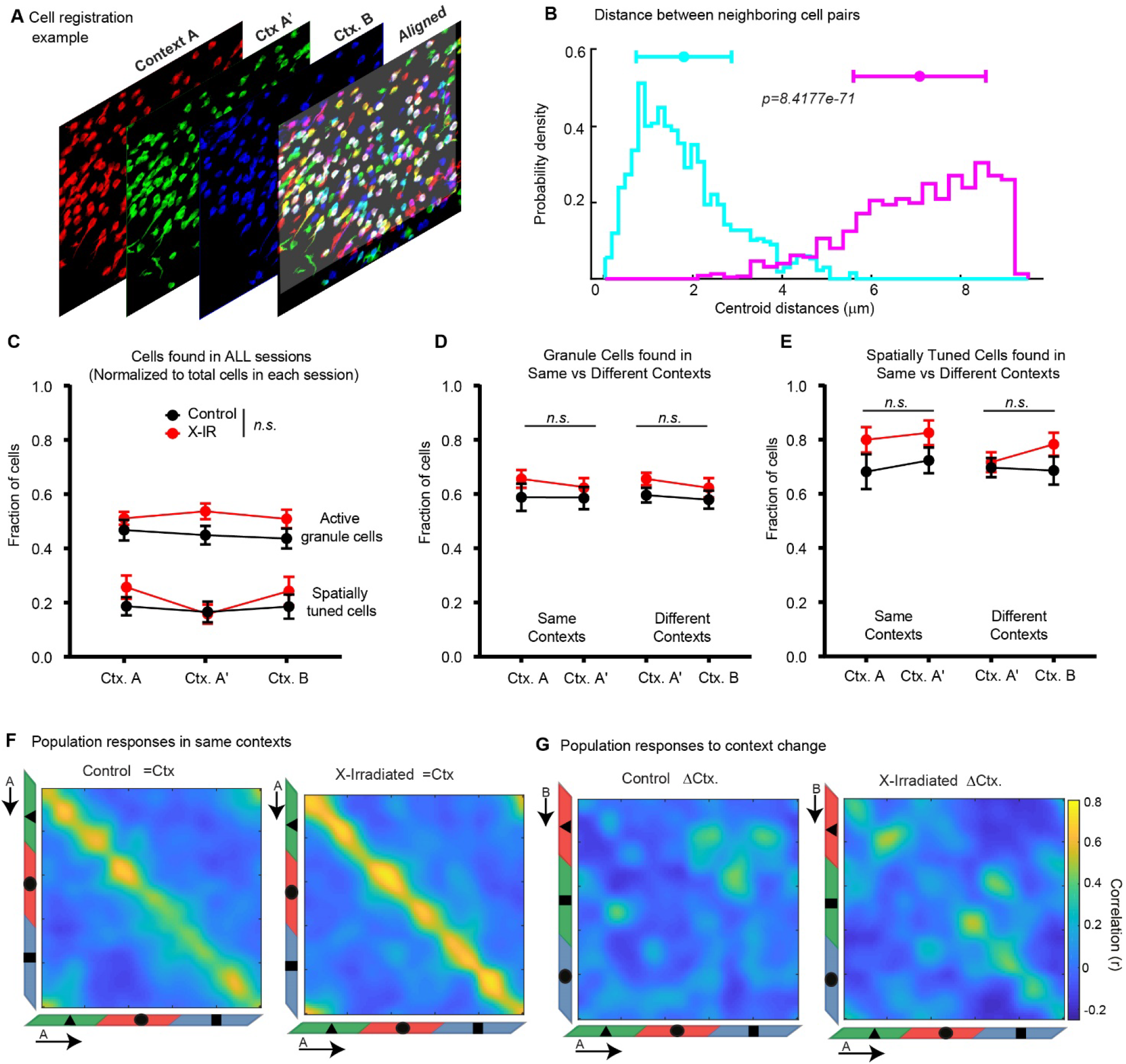
Chronic ablation of neurogenesis does not alter reactivation of same GC populations but changes remapping of spatial representations in response to contextual changes (related to Figure 1) **A)** Representative alignment of spatial footprints for cells segmented across sessions within a day in different context sessions in a single imaging field. **B)** Distribution of centroid distances between registered (blue) and non-registered (pink) neighboring cell pairs (p=8.4×10^−71^, Mann-Whittney test). **C)** Comparison of the fraction of active granule cells (with at least 1 transient during a 15 min. session) or spatially tuned cells that are found in all three context sessions, normalized to all cells found in the same imaging field in control and X-IR mice. Active granule cells: *F*^2,45^ = 0.16, *P* = 0.84; Context *F*^2,45^ = 0.06, *P*=0.94; Genotype *F*^1,45^ = 3.95, *P* = 0.053; Spatially tuned granule cells: *F*^2,45^ = 0.52, *P* = 0.60; Context *F*^2,45^ = 0.29, *P*=0.94; Genotype *F*^1,45^ = 1.41, *P* = 0.24; 2-way ANOVA. **D)** The fraction of active granule cells that are also found (registered) in the same context sessions (left) or different context sessions (right) normalized to all cells found in the same imaging field in control and X-IR mice. Same Context (A-A’): Interaction *F*^1,30^ = 0.13, *P* = 0.72; Context *F*^1,30^ = 0.18, *P*=0.66; Genotype *F*^1,30^ = 1.64, *P* = 0.21; Different Context (A’-B): Interaction *F*^1,30^ = 0.07, *P* = 0.79; Context *F*^1,30^ = 0.66, *P*=0.42; Genotype *F*^1,30^ = 2.95, *P* = 0.09; 2-way ANOVA. **E)** The fraction of spatially tuned granule cells that are also found (registered) in the same context sessions (left) or different context sessions (right) normalized to the active granule cells in the same imaging field in control and X-IR mice. Same Context (A-A’): Interaction *F*^1,30^ = 0.12, *P* = 0.73; Context *F*^1,30^ = 0.20, *P*=0.65; Genotype *F*^1,30^ = 3.60, *P* = 0.06; Different Context (A’-B): Interaction *F*^1,30^ = 0.82, *P* = 0.37; Context *F*^1,30^ = 0.43, *P*=0.52; Genotype *F*^1,30^ = 1.89, *P* = 0.18; 2-way ANOVA. Note that the comparisons in B-E test the stability of imaging fields in controls and X-IR mice that may affect selection bias during the registration algorithm for cells registered in other sessions regardless of their spatial tuning profiles, for comparisons of reactivation of spatially tuned ensembles in same vs different contexts see Supplementary Figures 3B, C, E, F. **F)** Population vector (PV) correlations of granule cells with significant fields in same contexts at each treadmill position in control (left) and X-IR (right) mice. For calculating PVs, lap-averaged spatial firing rate maps of all tuned cells in the second session of Context A (A’) were correlated with that of the first session of Context A. **G)** Population vector (PV) correlations of granule cells with significant fields in different contexts at each treadmill position in control (left) and X-IR (right) mice. Lap-averaged spatial firing rate maps of all tuned cells in the second session of Context A were correlated with that of the Context B.

**Supplementary Figure 2:**
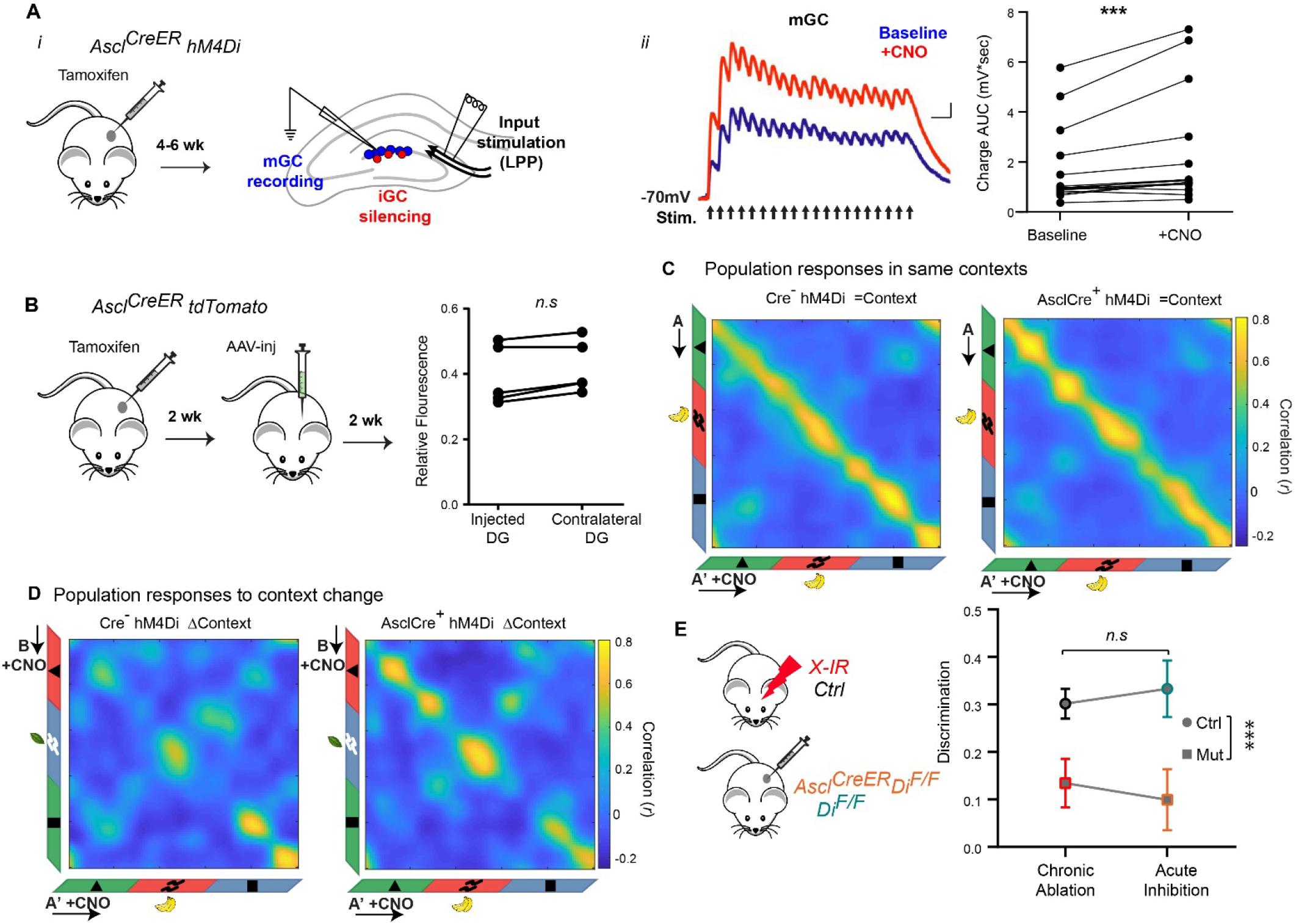
Acute silencing of 4-week-old iGCs (related to Figure 2). **A)** (*i*) Ascl^CreER^;Di^F/F^ mice were injected with TAM 4-6 weeks later acute slice were prepared (left). A bipolar stimulating electrode was positioned in the lateral perforant path (LPP, the outermost section of the DG molecular layer). Whole cell current clamp recordings were performed on mGCs before (baseline) and during bath application of CNO (5mM). (*ii*) Average evoked synaptic potentials in a representative mGC during baseline (blue) and CNO application (red). AUC indicates area above baseline (depolarization). *W* = 112, \**P*^*Baseline-CNO*^= 0.0004, Wilcoxon matched pairs signed rank test, N=15 cells from 8 mice. **B)** Ascl^CreER^;TdTomato^F/F^ mice were injected with TAM followed by unilateral AAV-GCaMp7s injection into the dorsal DG 2 weeks later (left). Quantification of TdTomato signal in upper blade of the DG shown in Figure 2B. *W* = 13, *P*^*IR-Day – IR-Week*^ = 0.125, Wilcoxon matched pairs signed rank test. N=5 mice, 3 matched sections/mouse. **C)** Population vector (PV) correlations of granule cells with significant fields in same contexts at each treadmill position in control (left) and X-IR (right) mice. For calculating PVs, lap-averaged spatial firing rate maps of all tuned cells in the second session of Context A (A’) were correlated with that of the first session of Context A. **D)** Population vector (PV) correlations of granule cells with significant fields in different contexts at each treadmill position in control (left) and X-IR (right) mice. Lap-averaged spatial firing rate maps of all tuned cells in the second session of Context A were correlated with that of the Context B. **E)** The effect size of Control - X-IR (Cohen’s *d*=1.48, *r*=0.59) and *Di* ^F/F^ - *Ascl*^*CreER*^ *Di* ^F/F^ (Cohen’s *d*=1.37, *r*=0.56) cohorts were comparable. Further, comparison of the context discrimination index calculated as: (Error ^Different Context^ - Error ^Same Context^)/ (Error ^Different Context^ + Error ^Same Context^), in both groups of mice. Interaction *F*^1,26^ = 0.12, *P* = 0.089; Manipulation *F*^1,26^ = 0.008, *P*=0.92; Genotype *F*^1,26^ = 112.6, \*\*\**P* < 0.0001; 2-way ANOVA.

**Supplementary Figure 3:**
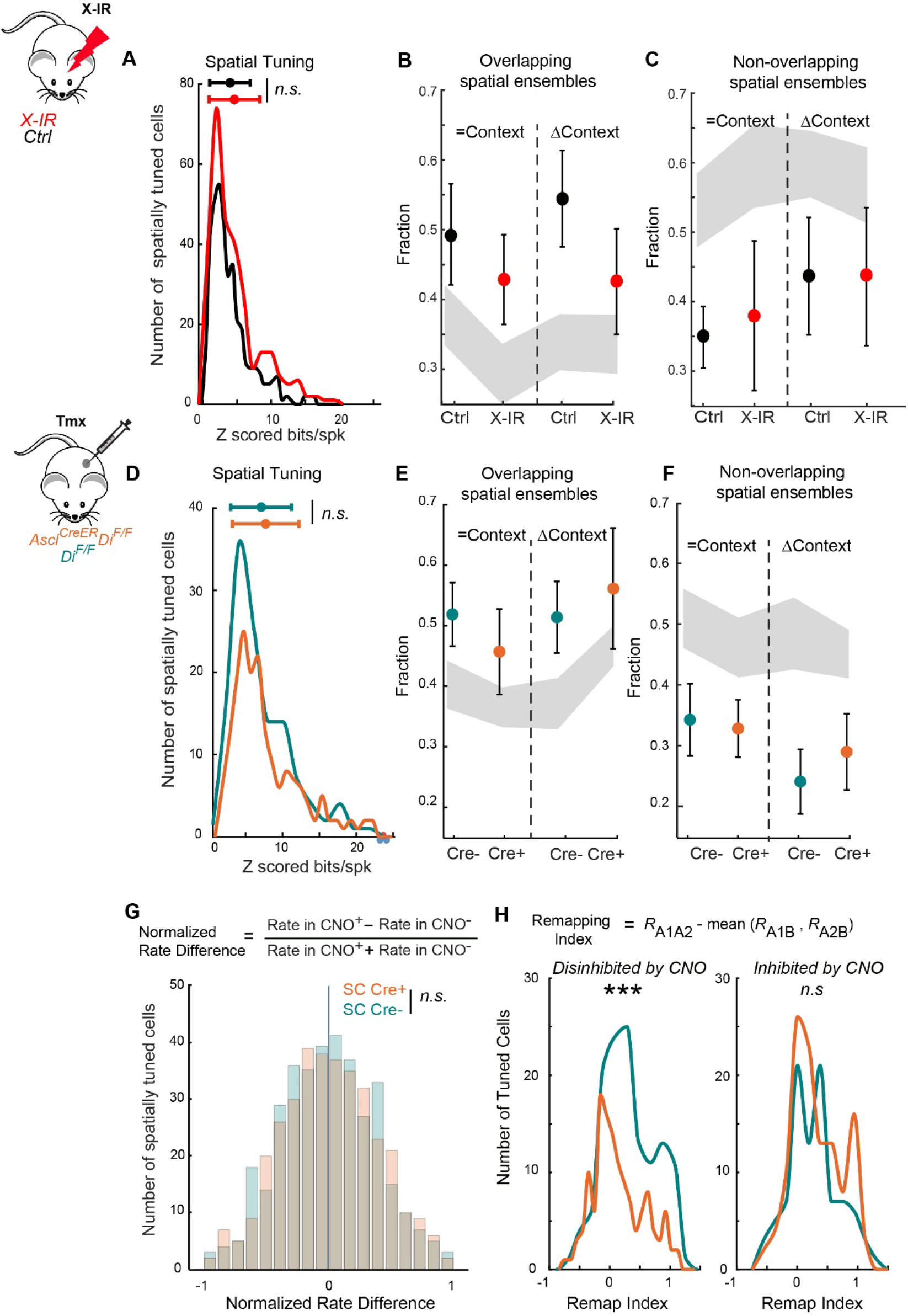
Global activity profiles of mature granule cells as a result of chronic ablation or transient inhibition of iGCs (related to Figure 3). **A)** The distribution of the spatial information contents of tuned granule cells in control and X-IR mice. *P* = 0.088, Control: 3.96 ± 0.15, X-IR: 4.60 ± 0.22 for granule cells that are significantly tuned in at least one session of the context discrimination task. **B)** Fraction of cross-session registered cells found in Context A’ that encoded space during exposure to same or different contexts in control and X-IR mice. χ^2^=2.46, p=0.48. **C)** Fraction of cross-session registered cells found in Conext A’ that did not encode space during exposure to same or different contexts in control and X-IR mice. χ^2^=1.52, p=0.67. **D)** The distribution of the spatial information contents of tuned granule cells in Di^F/F^ controls and Ascl^CreER^;Di^F/F^ mice. *P* = 0.56, Di^F/F^: 7.10 ± 0.37, Ascl^CreER^;Di^F/F^: 7.71 ± 0.34 for granule cells that are significantly tuned in both baseline and CNO injected sessions of the context discrimination task. **E)** Fraction of cross-session registered cells found in Context A’ that encoded space during exposure to same or different contexts in Di^F/F^ controls and Ascl^CreER^;Di^F/F^ mice. χ^2^=2.79, p=0.42. **F)** Fraction of cross-session registered cells found in Conext A’ that did not encode space during exposure to same or different contexts in Di^F/F^ and Ascl^CreER^;Di^F/F^ mice. χ^2^=1.27, p=0.73 **G)** Distribution of GCs’ CNO induced normalized rate differences in Ascl^CreER^;Di^F/F^ mice compared to those in Di^F/F^ controls. *P* = 0.29, Di^F/F^: 0.046 ± 0.017, Ascl^CreER^;Di^F/F^: 0.0413 ± 0.0172 **H)** The distribution of the context selectivity of spatially tuned cells that are disinhibited (right, \*\**P* = 0.002) or inhibited (left, *P*=0.39) by CNO in Ascl^CreER^;Di^F/F^ mice compared to those in Di^F/F^ controls. Gray areas represent 2.5^th^ and 97.5^th^% of null distributions for each cell type in B, C, E, F The null distributions are generated for each cell type by randomly permuting cell IDs of all cross-registered neurons (5,000 shufflings) and determining overlap among cell types. A-D,G,H: Two-sample Kolmogorov-Smirnov test. B-C-E-F: Kruskal Wallis test. Error bars, ± sem.

**Supplementary Figure 4.**
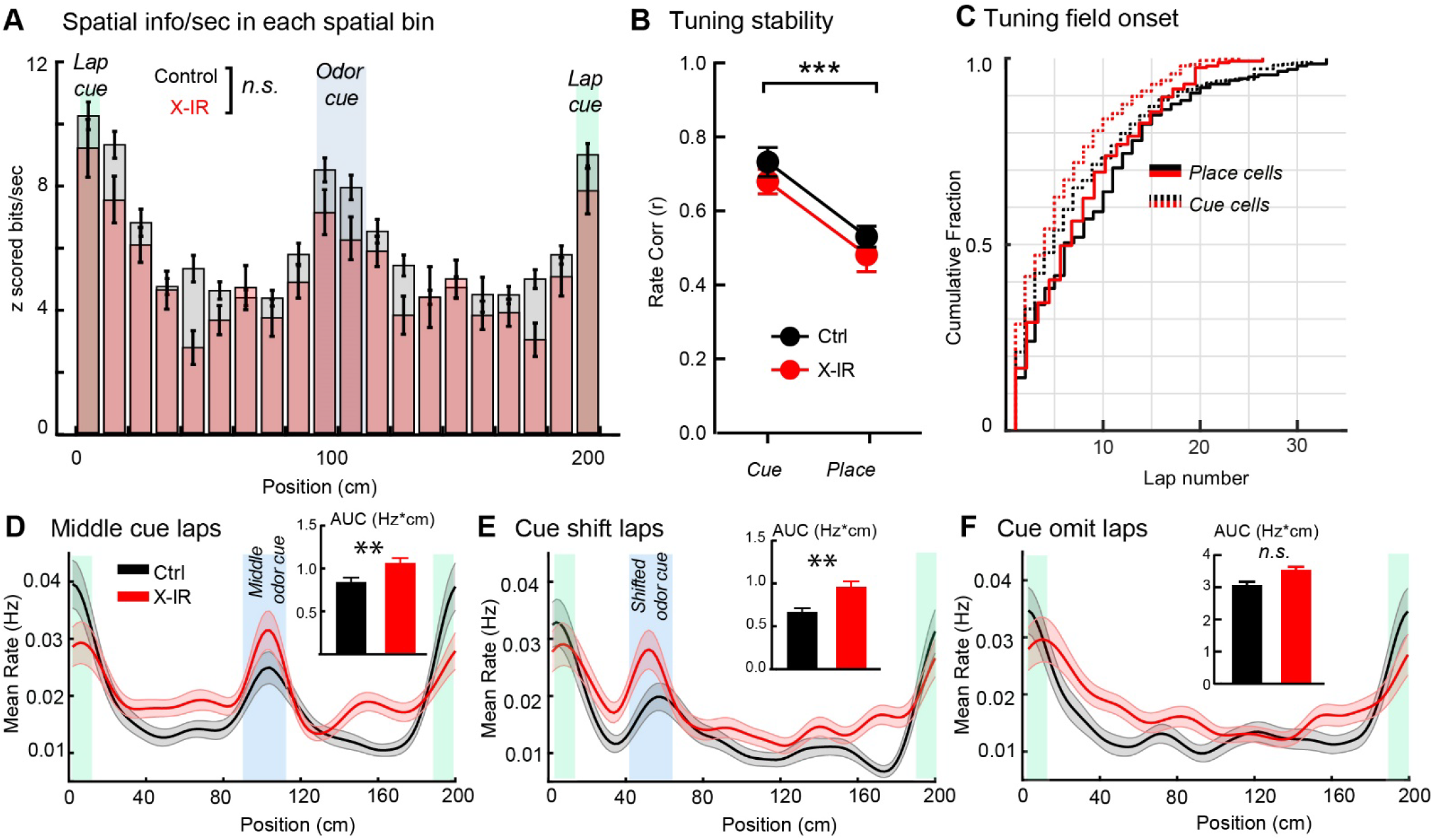
Effects of chronic ablation of neurogenesis on the functional segregation of cue and place responses in the DG (related to Figure 4). **A)** Mean spatial information by position showing similar enrichment of neurons with high spatial information around the sensory cues in both X-IR (pink) and control (gray) mice. Average Z scored spatial information for cells binned by tuning position during the spatial cue task performed by groups of mice. P=0.0926, Two-sample Kolmogorov-Smirnov test. Error bars, ± sem. **B)** Tuning consistency of cue and place cells in control and X-IR mice during spatial cue task. Firing rate correlation between first and last halves of the session. Red dotted lines in violin plots show median, black dotted lines show quartiles. Interaction *F*^1,24^ = 0.002, *P* = 0.96; Cell Type *F*^1,24^ = 29.12, \*\*\**P* < 0.0001; Genotype *F*^1,24^ = 1.876, *P*=0.1834; 2-way ANOVA. **C)** Emergence of cue and place responses. Cumulative distribution of spatial field onset lap for cue and place cells in control and X-IR mice during spatial cue task. Interaction *F*^1,617^ = 0.006, *P* = 0.94; Cell Type *F*^1,617^ = 2.43, *P* =0.12; Genotype *F*^1,617^ = 6.004, \**P*=0.014; 2-way ANOVA. **D)** Average spatial firing rates from all cells in Figure 4B during normal middle cue laps in control (black) and X-IR mice. (Inset) averaged area under the firing rate curves (AUC, Hz*cm) within the middle region corresponding to odor delivery during normal laps in control (black bar) and X-IR mice (red bar). Locations of the odor and lap cue are shown blue and green shaded areas, respectively. \*\**P*^*Ctrl-IR*^ = 0.0036. **E)** Average spatial firing rate by position for neurons on cue-shifted laps. (Inset) averaged area under the firing rate curves (AUC, Hz*cm) within the cue region during cue-shifted laps. \*\**P*^*Ctrl-IR*^ =0.002. **F)** Average spatial firing rate by position for neurons on cue-omitted laps. (Inset) averaged area under the firing rate curves (AUC, Hz*cm) during cue omitted laps. *P*^*Ctrl-IR*^ =0.055. D-F: Mann-Whitney test. Error bars, ± sem. **G)** Population vector (PV) correlations of granule cells with significant field at each treadmill position within a day in control (left) and X-IR (right) mice. For calculating PVs, lap-averaged spatial firing rate maps from normal middle cue laps of all tuned cells in Day1 were correlated with that of the second session from the dame day. **H)** Population vector (PV) correlations of granule cells at each treadmill position within a week in control (left) and X-IR (right) mice. Lap-averaged spatial firing rate maps from normal middle cue laps of all tuned cells from the first session were correlated with that of the session imaged 7 days later.

**Supplementary Figure 5:**
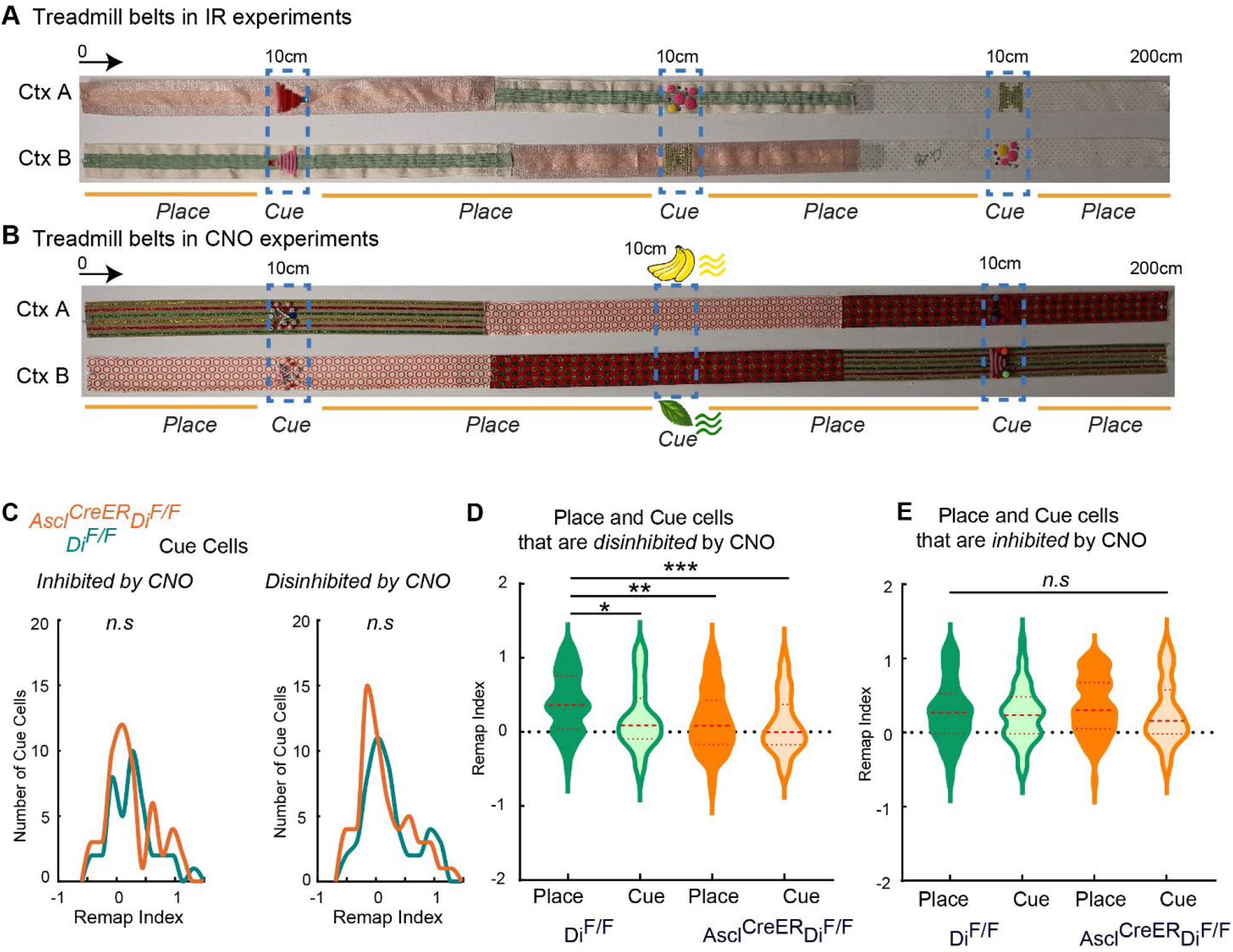
Effects of acute silencing of iGCs on the context selectivity of place and cue cells (related to Figure 5). **A)** Images of the two belts (A and B) used in the IR experiments. The order of the fabrics was changed and the three cues were affixed in a different arrangement. Cue encoding mGCs were defined as those with peak Ca^2+^ event rate within the boundaries of the affixed tactile cues (10 cm wide) while the place encoding mGCs were defined as those with peak Ca^2+^ event rate outside of the 10 cm boundary of the affixed tactile cues on the treadmill belt during the second exposure to context A. Direction of movement and the selected locations for the start of the belts are indicated. **B)** Images of the two belts (A and B) used in the CNO experiments. The order of the fabrics was changed, different odor cues were presented at one location (1 sec pulse) and the two cues were affixed in a different arrangement. Cue encoding mGCs were defined as those with peak Ca^2+^ event rate within the boundaries of the affixed tactile cues or the odor cue while the place encoding mGCs were defined as those with peak Ca^2+^ event rate outside of the cue locations on the treadmill belt during the second exposure to context A. Direction of movement and the selected locations for the start of the belts are indicated. **C)** The distribution of the remapping index of cue cells that are inhibited (left, *P*=0.63) or disinhibited (right, *P* = 0.28 or by CNO in Ascl^CreER^;Di^F/F^ mice compared to those in Di^F/F^ controls. Two-sample Kolmogorov-Smirnov test. **D)** Comparisons of the remapping indices of place and cue cells that are disinhibited by CNO in Di^F/F^ and Ascl^CreER^;Di^F/F^ mice. χ^2^=21.59, \*\*\**P* < 0.0001; \**P* ^*Cre-Place – Cre-Cue*^ = 0.039, \*\**P* ^*Cre-Place – Cre+ Place*^ = 0.0015, \*\*\**P* ^*Cre-Place – Cre+ Cue*^ = 0.0002, Kruskal Wallis test, Dunn’s multiple comparisons test. Dashed lines in violin plots show median, dotted lines show quartiles. **E)** Comparisons of the remapping indices of place and cue cells that are inhibited by CNO in Di^F/F^ and Ascl^CreER^;Di^F/F^ mice. χ^2^=2.7, *P* = 0.43; Kruskal Wallis test.

**Supplementary Figure 6:**
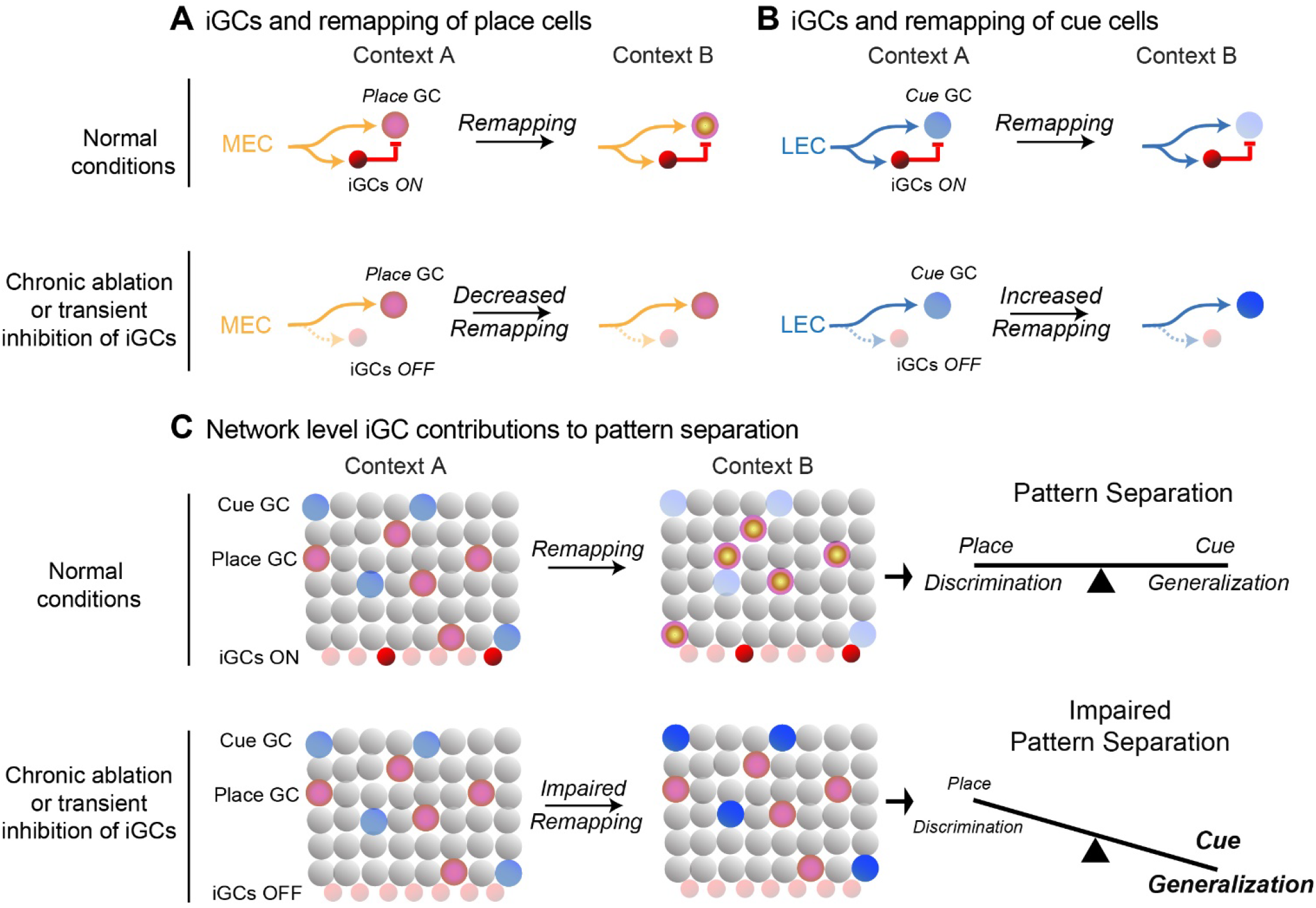
Model illustrating the contribution of adult born immature granule cells to pattern separation. **A)** (Top) Under normal conditions, place encoding mature granule cells represent self-motion information between different cues context A (yellow circle with pink center) and represent self-motion information in context B by decorrelating their firing fields (yellow circle with pink surround). This is likely mediated by the modulation of spatial information arriving from the medial entorhinal cortex (MEC, orange arrows). (Bottom) Loss of iGCs prevent the modulation of these inputs to the place GCs, resulting in decreased remapping place GCs between context A and B (yellow circles with pink center). **B)** (Top) Under normal conditions, cue encoding mature granule cells represent various sensory features of context A (light blue circle) and represent the same sensory cues in context B by modulating in their activity levels (lighter blue circle). This is likely mediated by the inhibitory modulation of sensory cue information arriving from the lateral entorhinal cortex (LEC, blue arrows) by immature adult born granule cells (iGCs, red circles). (Bottom) Chronic ablation of transient inhibition of iGCs prevent the inhibitory control of LEC inputs on cue GCs, resulting in high activity and increased remapping cue GCs between context A (light blue circle) and B (dark blue circle) even though the cues are at different locations of the sensory cues in different contexts. **C)** Representations of contexts A and B are made up of mature GCs encoding sensory cues and place information. Under normal conditions, pattern separation of contexts A and B occurs by rate remapping of cue cells to form distinct traces for same objects encountered in different locations and global remapping of place cells to form non-overlapping representations about animal’s location within different contexts. The balanced representation of sensory cue and self-motion information allows the network flexibly discriminate similar contexts and generalize common sensory features. Chronic ablation of transient inhibition of iGCs, impairs pattern separation by increasing remapping of sensory cue representations and decreasing the remapping of place cells and self-motion representations. In turn, this shifts the DG network state towards higher levels of sensory cue representation and less flexible self-motion representation resulting in over-generalization and lower discrimination.

## References

Anacker C, Luna VM, Stevens GS, Millette A, Shores R, Jimenez JC, Chen B, Hen R (2018) Hippocampal neurogenesis confers stress resilience by inhibiting the ventral dentate gyrus. Nature 559:98–102.

Belzung C, Willner P, Philippot P (2015) Depression: from psychopathology to pathophysiology. Curr Opin Neurobiol 30:24–30.

Burgess N, O’Keefe J (1996) Neuronal computations underlying the firing of place cells and their role in navigation. Hippocampus 6:749–762.

Burghardt NS, Park EH, Hen R, Fenton AA (2012) Adult-born hippocampal neurons promote cognitive flexibility in mice. Hippocampus 22:1795–1808.

Colgin LL, Moser EI, Moser M-B (2008) Understanding memory through hippocampal remapping. Trends Neurosci 31:469–477.

Danielson NB, Kaifosh P, Zaremba JD, Lovett-Barron M, Tsai J, Denny CA, Balough EM, Goldberg AR, Drew LJ, Hen R, Losonczy A, Kheirbek MA (2016) Distinct Contribution of Adult-Born Hippocampal Granule Cells to Context Encoding. Neuron 90:101–112.

Danielson NB, Turi GF, Ladow M, Chavlis S, Petrantonakis PC, Poirazi P, Losonczy A (2017) In Vivo Imaging of Dentate Gyrus Mossy Cells in Behaving Mice. Neuron 93:552-559.e4.

Denny CA, Burghardt NS, Schachter DM, Hen R, Drew MR (2012) 4-to 6-week-old adult-born hippocampal neurons influence novelty-evoked exploration and contextual fear conditioning. Hippocampus 22:1188–1201.

Diamantaki M, Frey M, Berens P, Preston-Ferrer P, Burgalossi A (2016) Sparse activity of identified dentate granule cells during spatial exploration. eLife 5.

Drew LJ, Kheirbek MA, Luna VM, Denny CA, Cloidt MA, Wu MV, Jain S, Scharfman HE, Hen R (2016) Activation of local inhibitory circuits in the dentate gyrus by adult-born neurons. Hippocampus 26:763–778.

Farrell JS, Lovett-Barron M, Klein PM, Sparks FT, Gschwind T, Ortiz AL, Ahanonu B, Bradbury S, Terada S, Oijala M, Hwaun E, Dudok B, Szabo G, Schnitzer MJ, Deisseroth K, Losonczy A, Soltesz I (2021) Supramammillary regulation of locomotion and hippocampal activity. Science 374:1492– 1496.

Fyhn M, Molden S, Witter MP, Moser EI, Moser M-B (2004) Spatial representation in the entorhinal cortex. Science 305:1258–1264.

Ge S, Yang C-H, Hsu K-S, Ming G-L, Song H (2007) A critical period for enhanced synaptic plasticity in newly generated neurons of the adult brain. Neuron 54:559–566.

Gómez-Ocádiz R, Trippa M, Zhang C-L, Posani L, Cocco S, Monasson R, Schmidt-Hieber C (2022) A synaptic signal for novelty processing in the hippocampus. Nat Commun 13:4122.

GoodSmith D, Chen X, Wang C, Kim SH, Song H, Burgalossi A, Christian KM, Knierim JJ (2017) Spatial Representations of Granule Cells and Mossy Cells of the Dentate Gyrus. Neuron 93:677-690.e5.

GoodSmith D, Kim SH, Puliyadi V, Ming G, Song H, Knierim JJ, Christian KM (2022) Flexible encoding of objects and space in single cells of the dentate gyrus. Curr Biol 32:1088-1101.e5.

Grosmark AD, Buzsáki G (2016) Diversity in neural firing dynamics supports both rigid and learned hippocampal sequences. Science 351:1440–1443.

Grosmark AD, Sparks FT, Davis MJ, Losonczy A (2021) Reactivation predicts the consolidation of unbiased long-term cognitive maps. Nat Neurosci 24:1574–1585.

Gu Y, Arruda-Carvalho M, Wang J, Janoschka SR, Josselyn SA, Frankland PW, Ge S (2012) Optical controlling reveals time-dependent roles for adult-born dentate granule cells. Nat Neurosci 15:1700–1706.

Hainmueller T, Bartos M (2018) Parallel emergence of stable and dynamic memory engrams in the hippocampus. Nature 558:292–296.

Hargreaves EL, Rao G, Lee I, Knierim JJ (2005) Major dissociation between medial and lateral entorhinal input to dorsal hippocampus. Science 308:1792–1794.

Huckleberry KA, Shue F, Copeland T, Chitwood RA, Yin W, Drew MR (2018) Dorsal and ventral hippocampal adult-born neurons contribute to context fear memory. Neuropsychopharmacol Off Publ Am Coll Neuropsychopharmacol 43:2487–2496.

Ikrar T, Guo N, He K, Besnard A, Levinson S, Hill A, Lee H-K, Hen R, Xu X, Sahay A (2013) Adult neurogenesis modifies excitability of the dentate gyrus. Front Neural Circuits 7:204.

Johnston S, Parylak SL, Kim S, Mac N, Lim C, Gallina I, Bloyd C, Newberry A, Saavedra CD, Novak O, Gonçalves JT, Gage FH, Shtrahman M (2021) AAV ablates neurogenesis in the adult murine hippocampus Lucassen P, Colgin LL, eds. eLife 10:e59291.

Johnston ST, Shtrahman M, Parylak S, Gonçalves JT, Gage FH (2016) Paradox of pattern separation and adult neurogenesis: A dual role for new neurons balancing memory resolution and robustness. Neurobiol Learn Mem 129:60–68.

Jung D, Kim S, Sariev A, Sharif F, Kim D, Royer S (2019) Dentate granule and mossy cells exhibit distinct spatiotemporal responses to local change in a one-dimensional landscape of visual-tactile cues. Sci Rep 9:9545.

Kheirbek MA, Klemenhagen KC, Sahay A, Hen R (2012) Neurogenesis and generalization: a new approach to stratify and treat anxiety disorders. Nat Neurosci 15:1613–1620.

Kim EJ, Ables JL, Dickel LK, Eisch AJ, Johnson JE (2011) Ascl1 (Mash1) Defines Cells with Long-Term Neurogenic Potential in Subgranular and Subventricular Zones in Adult Mouse Brain. PLOS ONE 6:e18472.

Knierim JJ, Neunuebel JP (2016) Tracking the flow of hippocampal computation: Pattern separation, pattern completion, and attractor dynamics. Neurobiol Learn Mem 129:38–49.

Knierim JJ, Neunuebel JP, Deshmukh SS (2014) Functional correlates of the lateral and medial entorhinal cortex: objects, path integration and local-global reference frames. Philos Trans R Soc Lond B Biol Sci 369:20130369.

Kubie JL, Levy ERJ, Fenton AA (2020) Is hippocampal remapping the physiological basis for context? Hippocampus 30:851–864.

Lacefield CO, Itskov V, Reardon T, Hen R, Gordon JA (2012) Effects of adult-generated granule cells on coordinated network activity in the dentate gyrus. Hippocampus 22:106–116.

Lange I, Goossens L, Michielse S, Bakker J, Lissek S, Papalini S, Verhagen S, Leibold N, Marcelis M, Wichers M, Lieverse R, van Os J, van Amelsvoort T, Schruers K (2017) Behavioral pattern separation and its link to the neural mechanisms of fear generalization. Soc Cogn Affect Neurosci 12:1720–1729.

Latuske P, Kornienko O, Kohler L, Allen K (2017) Hippocampal Remapping and Its Entorhinal Origin. Front Behav Neurosci 11:253.

Li Y-D, Luo Y-J, Chen Z-K, Quintanilla L, Cherasse Y, Zhang L, Lazarus M, Huang Z-L, Song J (2022) Hypothalamic modulation of adult hippocampal neurogenesis in mice confers activity-dependent regulation of memory and anxiety-like behavior. Nat Neurosci 25:630–645.

Lissek S, Rabin S, Heller RE, Lukenbaugh D, Geraci M, Pine DS, Grillon C (2010) Overgeneralization of conditioned fear as a pathogenic marker of panic disorder. Am J Psychiatry 167:47–55.

Lu L, Leutgeb JK, Tsao A, Henriksen EJ, Leutgeb S, Barnes CA, Witter MP, Moser M-B, Moser EI (2013) Impaired hippocampal rate coding after lesions of the lateral entorhinal cortex. Nat Neurosci 16:1085–1093.

Luna VM, Anacker C, Burghardt NS, Khandaker H, Andreu V, Millette A, Leary P, Ravenelle R, Jimenez JC, Mastrodonato A, Denny CA, Fenton AA, Scharfman HE, Hen R (2019) Adult-born hippocampal neurons bidirectionally modulate entorhinal inputs into the dentate gyrus. Science 364:578–583.

Madisen L, Zwingman TA, Sunkin SM, Oh SW, Zariwala HA, Gu H, Ng LL, Palmiter RD, Hawrylycz MJ, Jones AR, Lein ES, Zeng H (2010) A robust and high-throughput Cre reporting and characterization system for the whole mouse brain. Nat Neurosci 13:133–140.

Marín-Burgin A, Mongiat LA, Pardi MB, Schinder AF (2012) Unique processing during a period of high excitation/inhibition balance in adult-born neurons. Science 335:1238–1242.

Marr D (1971) Simple memory: a theory for archicortex. Philos Trans R Soc Lond B Biol Sci 262:23–81.

McAvoy K, Besnard A, Sahay A (2015) Adult hippocampal neurogenesis and pattern separation in DG: a role for feedback inhibition in modulating sparseness to govern population-based coding. Front Syst Neurosci 9:120.

McHugh SB, Lopes-dos-Santos V, Gava GP, Hartwich K, Tam SKE, Bannerman DM, Dupret D (2022) Adult-born dentate granule cells promote hippocampal population sparsity. Nat Neurosci:1–11.

Miao C, Cao Q, Ito HT, Yamahachi H, Witter MP, Moser M-B, Moser EI (2015) Hippocampal Remapping after Partial Inactivation of the Medial Entorhinal Cortex. Neuron 88:590–603.

Nakashiba T, Cushman JD, Pelkey KA, Renaudineau S, Buhl DL, McHugh TJ, Rodriguez Barrera V, Chittajallu R, Iwamoto KS, McBain CJ, Fanselow MS, Tonegawa S (2012) Young dentate granule cells mediate pattern separation, whereas old granule cells facilitate pattern completion. Cell 149:188–201.

Ogando MB, Pedroncini O, Federman N, Romano SA, Brum LA, Lanuza GM, Refojo D, Marin-Burgin A (2021) Cholinergic modulation of dentate gyrus processing through dynamic reconfiguration of inhibitory circuits. Cell Rep 36:109572.

O’Keefe J, Dostrovsky J (1971) The hippocampus as a spatial map. Preliminary evidence from unit activity in the freely-moving rat. Brain Res 34:171–175.

Pnevmatikakis EA, Giovannucci A (2017) NoRMCorre: An online algorithm for piecewise rigid motion correction of calcium imaging data. J Neurosci Methods 291:83–94.

Ray RS, Corcoran AE, Brust RD, Kim JC, Richerson GB, Nattie E, Dymecki SM (2011) Impaired respiratory and body temperature control upon acute serotonergic neuron inhibition. Science 333:637–642.

Rueckemann JW, DiMauro AJ, Rangel LM, Han X, Boyden ES, Eichenbaum H (2016) Transient optogenetic inactivation of the medial entorhinal cortex biases the active population of hippocampal neurons. Hippocampus 26:246–260.

Sahay A, Scobie KN, Hill AS, O’Carroll CM, Kheirbek MA, Burghardt NS, Fenton AA, Dranovsky A, Hen R (2011) Increasing adult hippocampal neurogenesis is sufficient to improve pattern separation. Nature 472:466–470.

Santarelli L, Saxe M, Gross C, Surget A, Battaglia F, Dulawa S, Weisstaub N, Lee J, Duman R, Arancio O, Belzung C, Hen R (2003) Requirement of Hippocampal Neurogenesis for the Behavioral Effects of Antidepressants. Science 301:805–809.

Schlesiger MI, Boublil BL, Hales JB, Leutgeb JK, Leutgeb S (2018) Hippocampal Global Remapping Can Occur without Input from the Medial Entorhinal Cortex. Cell Rep 22:3152–3159.

Schmidt-Hieber C, Jonas P, Bischofberger J (2004) Enhanced synaptic plasticity in newly generated granule cells of the adult hippocampus. Nature 429:184–187.

Senzai Y, Buzsáki G (2017) Physiological Properties and Behavioral Correlates of Hippocampal Granule Cells and Mossy Cells. Neuron 93:691-704.e5.

Sheffield MEJ, Adoff MD, Dombeck DA (2017) Increased Prevalence of Calcium Transients across the Dendritic Arbor during Place Field Formation. Neuron 96:490-504.e5.

Sheintuch L, Rubin A, Brande-Eilat N, Geva N, Sadeh N, Pinchasof O, Ziv Y (2017) Tracking the Same Neurons across Multiple Days in Ca2+ Imaging Data. Cell Rep 21:1102–1115.

Skaggs WE, McNaughton BL, Gothard KM (1993) An Information-Theoretic Approach to Deciphering the Hippocampal Code. In: Advances in Neural Information Processing Systems 5 (Hanson SJ, Cowan JD, Giles CL, eds), pp 1030–1037. Morgan-Kaufmann. Available at: http://papers.nips.cc/paper/671-an-information-theoretic-approach-to-deciphering-the-hippocampal-code.pdf [Accessed January 27, 2020].

Snyder JS, Kee N, Wojtowicz JM (2001) Effects of adult neurogenesis on synaptic plasticity in the rat dentate gyrus. J Neurophysiol 85:2423–2431.

Temprana SG, Mongiat LA, Yang SM, Trinchero MF, Alvarez DD, Kropff E, Giacomini D, Beltramone N, Lanuza GM, Schinder AF (2015) Delayed coupling to feedback inhibition during a critical period for the integration of adult-born granule cells. Neuron 85:116–130.

Tuncdemir SN, Grosmark AD, Turi GF, Shank A, Bowler JC, Ordek G, Losonczy A, Hen R, Lacefield CO (2022) Parallel processing of sensory cue and spatial information in the dentate gyrus. Cell Rep 38:110257.

Tuncdemir SN, Lacefield CO, Hen R (2019) Contributions of adult neurogenesis to dentate gyrus network activity and computations. Behav Brain Res 374:112112.

Vivar C, Potter MC, Choi J, Lee J-Y, Stringer TP, Callaway EM, Gage FH, Suh H, van Praag H (2012) Monosynaptic inputs to new neurons in the dentate gyrus. Nat Commun 3:1107.

Witter MP (2007) The perforant path: projections from the entorhinal cortex to the dentate gyrus. Prog Brain Res 163:43–61.

Woods NI, Stefanini F, Apodaca-Montano DL, Tan IMC, Biane JS, Kheirbek MA (2020) The Dentate Gyrus Classifies Cortical Representations of Learned Stimuli. Neuron 107:173-184.e6.

Woods NI, Vaaga CE, Chatzi C, Adelson JD, Collie MF, Perederiy JV, Tovar KR, Westbrook GL (2018) Preferential Targeting of Lateral Entorhinal Inputs onto Newly Integrated Granule Cells. J Neurosci Off J Soc Neurosci 38:5843–5853.

Yassa MA, Stark CEL (2011) Pattern separation in the hippocampus. Trends Neurosci 34:515–525.

Zaremba JD, Diamantopoulou A, Danielson NB, Grosmark AD, Kaifosh PW, Bowler JC, Liao Z, Sparks FT, Gogos JA, Losonczy A (2017) Impaired hippocampal place cell dynamics in a mouse model of the 22q11.2 deletion. Nat Neurosci 20:1612–1623.

