## Supplemental figures for "Adult born hippocampal granule cells promote pattern separation by bidirectionally modulating the remapping of place and cue cells"

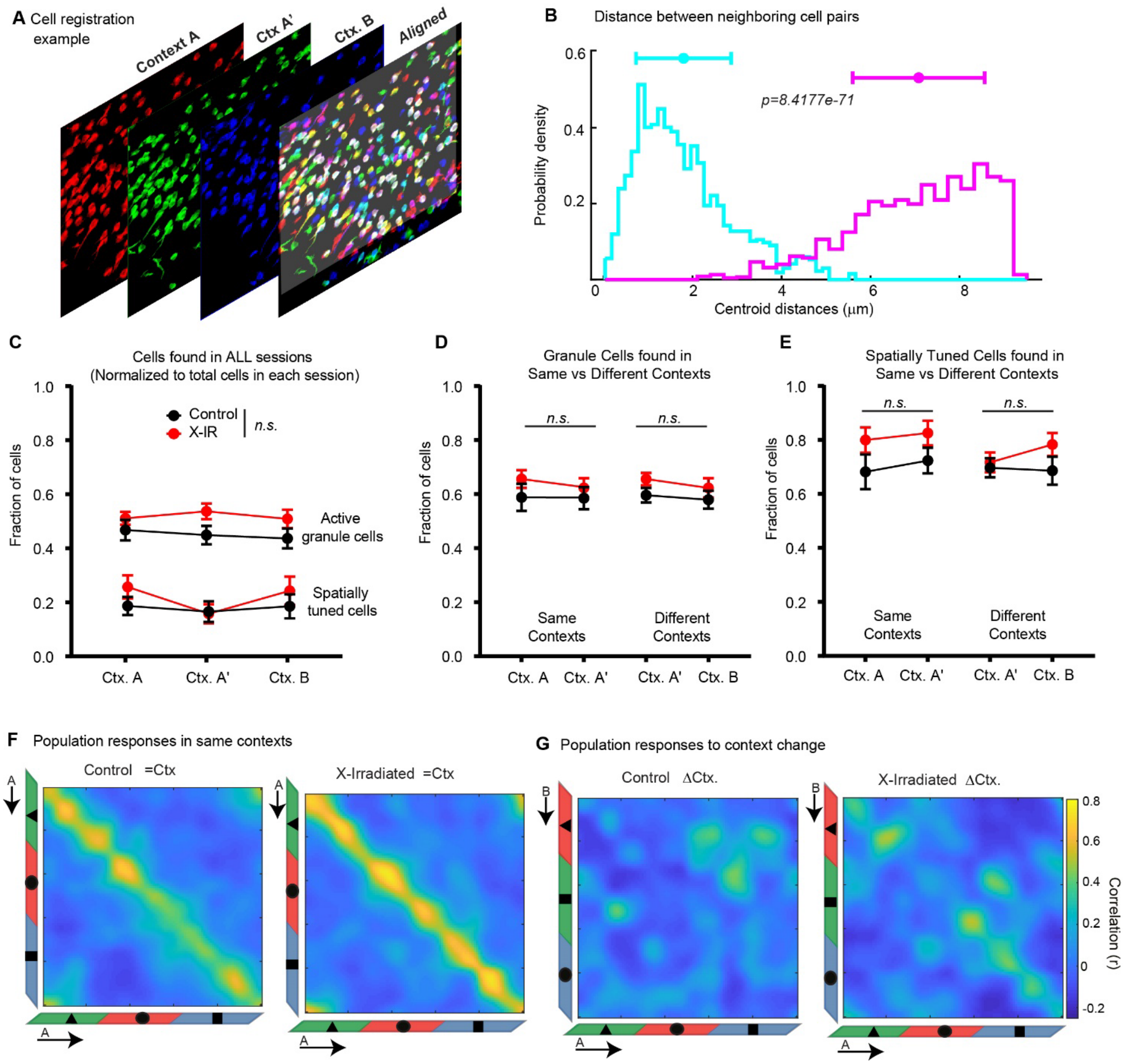


**Supplementary Figure 1. Chronic ablation of neurogenesis does not alter reactivation of same GC populations but changes remapping of spatial representations in response to contextual changes (related to Figure 1)**

**A)** Representative alignment of spatial footprints for cells segmented across sessions within a day in different context sessions in a single imaging field.

**B)** Distribution of centroid distances between registered (blue) and non-registered (pink) neighboring cell pairs (p=8.4x10$^{-71}$, Mann-Whittney test).

**C)** Comparison of the fraction of active granule cells (with at least 1 transient during a 15 min. session) or spatially tuned cells that are found in all three context sessions, normalized to all cells found in the same imaging field in control and X-IR mice. Active granule cells: $F_{2,45}$ = 0.16, $P$ = 0.84; Context $F_{2,45}$ = 0.06, $P$=0.94; Genotype $F_{1,45}$ = 3.95, $P$ = 0.053; Spatially tuned granule cells: $F_{2,45}$ = 0.52, $P$ = 0.60; Context $F_{2,45}$ = 0.29, $P$=0.94; Genotype $F_{1,45}$ = 1.41, $P$ = 0.24; 2-way ANOVA.

**D)** The fraction of active granule cells that are also found (registered) in the same context sessions (left) or different context sessions (right) normalized to all cells found in the same imaging field in control and X-IR mice. Same Context (A-A'): Interaction $F_{1,30}$ = 0.13, $P$ = 0.72; Context $F_{1,30}$ = 0.18, $P$=0.66;

Genotype $F_{1,30}$ = 1.64, $P$ = 0.21; Different Context (A'-B): Interaction $F_{1,30}$ = 0.07, $P$ = 0.79; Context $F_{1,30}$ = 0.66, $P$=0.42; Genotype $F_{1,30}$ = 2.95, $P$ = 0.09; 2-way ANOVA.

**E)** The fraction of spatially tuned granule cells that are also found (registered) in the same context sessions (left) or different context sessions (right) normalized to the active granule cells in the same imaging field in control and X-IR mice. Same Context (A-A'): Interaction $F_{1,30}$ = 0.12, $P$ = 0.73; Context $F_{1,30}$ = 0.20, $P$=0.65; Genotype $F_{1,30}$ = 3.60, $P$ = 0.06; Different Context (A'-B): Interaction $F_{1,30}$ = 0.82, $P$ = 0.37; Context $F_{1,30}$ = 0.43, $P$=0.52; Genotype $F_{1,30}$ = 1.89, $P$ = 0.18; 2-way ANOVA. Note that the comparisons in B-E test the stability of imaging fields in controls and X-IR mice that may affect selection bias during the registration algorithm for cells registered in other sessions regardless of their spatial tuning profiles, for comparisons of reactivation of spatially tuned ensembles in same vs different contexts see Supplementary Figures 3B, C, E, F.

**F)** Population vector (PV) correlations of granule cells with significant fields in same contexts at each treadmill position in control (left) and X-IR (right) mice. For calculating PVs, lap-averaged spatial firing rate maps of all tuned cells in the second session of Context A (A') were correlated with that of the first session of Context A.

**G)** Population vector (PV) correlations of granule cells with significant fields in different contexts at each treadmill position in control (left) and X-IR (right) mice. Lap-averaged spatial firing rate maps of all tuned cells in the second session of Context A were correlated with that of the Context B.

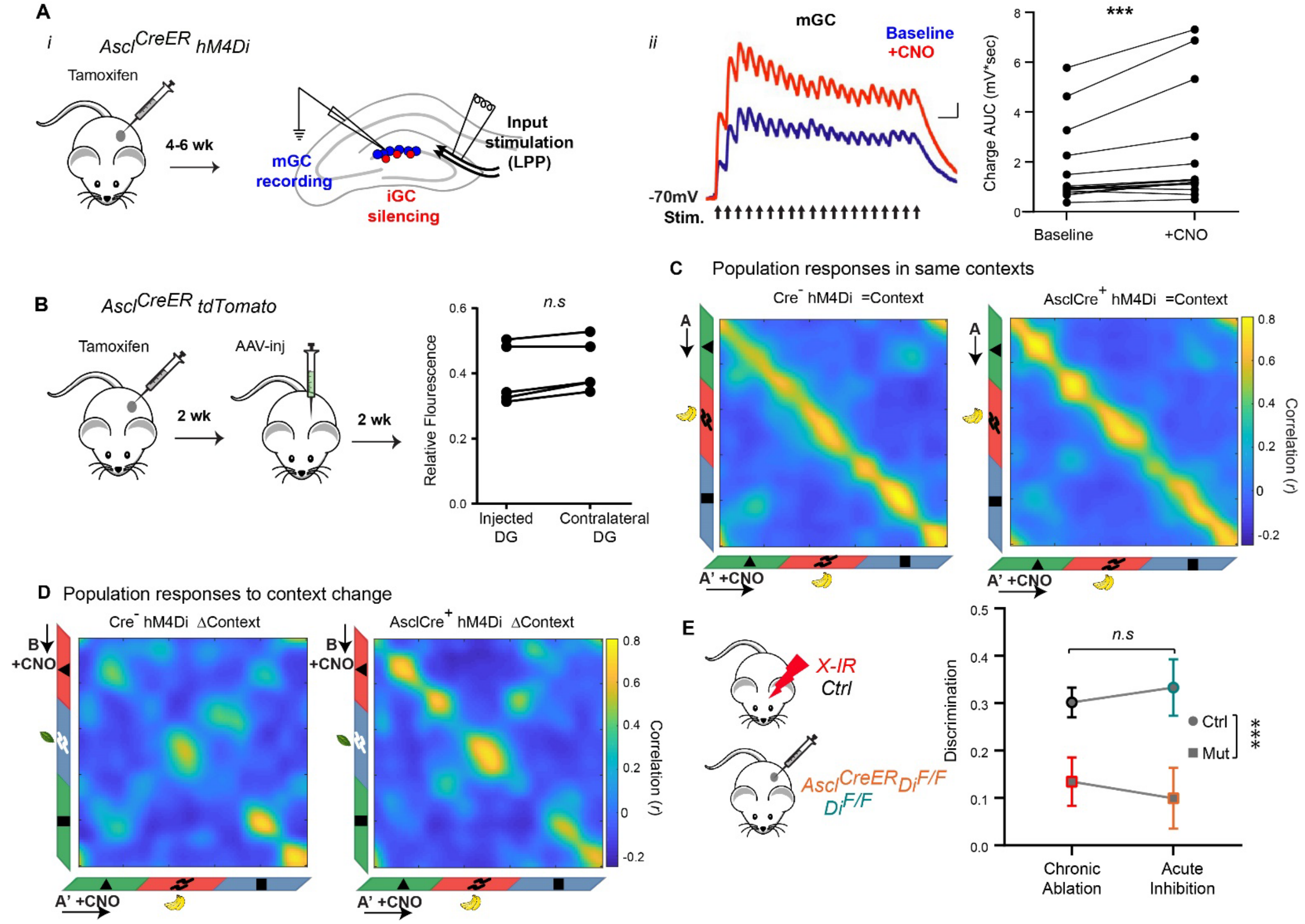


**Supplementary Figure 2: Acute silencing of 4-week-old iGCs (related to Figure 2).**

**A)** (*i*) $Ascl^{CreER}$;$Di^{F/F}$ mice were injected with TAM 4-6 weeks later acute slice were prepared (left). A bipolar stimulating electrode was positioned in the lateral perforant path (LPP, the outermost section of the DG molecular layer). Whole cell current clamp recordings were performed on mGCs before (baseline) and during bath application of CNO (5mM). (*ii*) Average evoked synaptic potentials in a representative mGC during baseline (blue) and CNO application (red). AUC indicates area above baseline (depolarization). $W = 112$, $*P_{Baseline\text{-}CNO} = 0.0004$, Wilcoxon matched pairs signed rank test, N=15 cells from 8 mice.

**B)** $Ascl^{CreER}$;$TdTomato^{F/F}$ mice were injected with TAM followed by unilateral AAV-GCaMp7s injection into the dorsal DG 2 weeks later (left). Quantification of TdTomato signal in upper blade of the DG shown in Figure 2B. $W = 13$, $P_{IR\text{-}Day - IR\text{-}Week} = 0.125$, Wilcoxon matched pairs signed rank test. N=5 mice, 3 matched sections/mouse.

**C)** Population vector (PV) correlations of granule cells with significant fields in same contexts at each treadmill position in control (left) and X-IR (right) mice. For calculating PVs, lap-averaged spatial firing rate maps of all tuned cells in the second session of Context A (A') were correlated with that of the first session of Context A.

**D)** Population vector (PV) correlations of granule cells with significant fields in different contexts at each treadmill position in control (left) and X-IR (right) mice. Lap-averaged spatial firing rate maps of all tuned cells in the second session of Context A were correlated with that of the Context B.

**E)** The effect size of Control - X-IR (Cohen's $d$=1.48, $r$=0.59) and $Di^{F/F}$ - $Ascl^{CreER}$ $Di^{F/F}$ (Cohen's $d$=1.37, $r$=0.56) cohorts were comparable. Further, comparison of the context discrimination index calculated as: $(Error_{Different\ Context} - Error_{Same\ Context})/(Error_{Different\ Context} + Error_{Same\ Context})$, in both groups of mice. Interaction $F_{1,26} = 0.12$, $P = 0.089$; Manipulation $F_{1,26} = 0.008$, $P$=0.92; Genotype $F_{1,26} = 112.6$, $***P < 0.0001$; 2-way ANOVA.

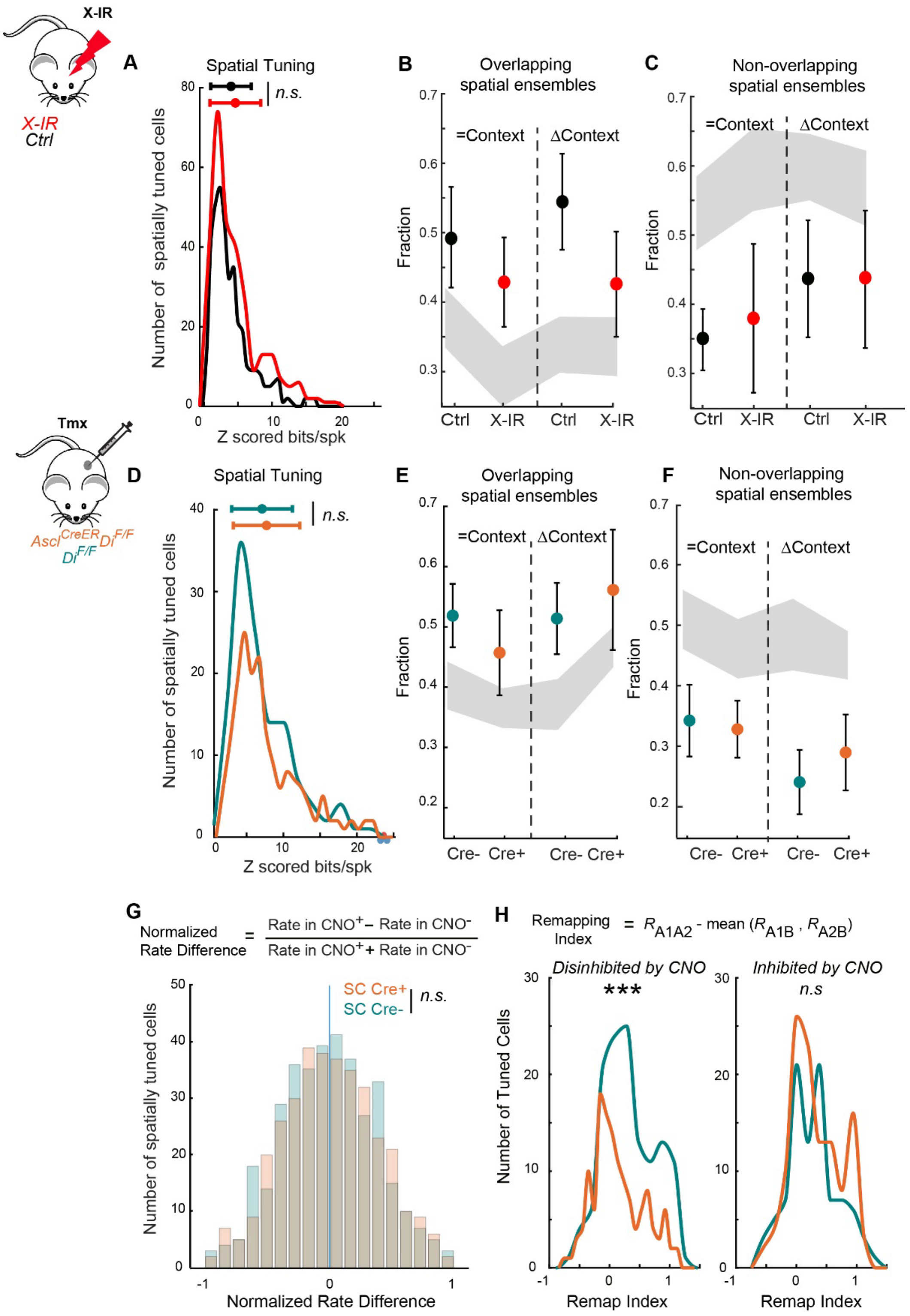


**Supplementary Figure 3: Global activity profiles of mature granule cells as a result of chronic ablation or transient inhibition of iGCs (related to Figure 3).**

- **A)** The distribution of the spatial information contents of tuned granule cells in control and X-IR mice. $P = 0.088$, Control: 3.96 ± 0.15, X-IR: 4.60 ± 0.22 for granule cells that are significantly tuned in at least one session of the context discrimination task.
- **B)** Fraction of cross-session registered cells found in Context A' that encoded space during exposure to same or different contexts in control and X-IR mice. $\chi^2=2.46$, $p=0.48$.

**C)** Fraction of cross-session registered cells found in Conext A' that did not encode space during exposure to same or different contexts in control and X-IR mice. $\chi^2$=1.52, p=0.67.

**D)** The distribution of the spatial information contents of tuned granule cells in $Di^{F/F}$ controls and $Ascl^{CreER}$;$Di^{F/F}$ mice. $P$ = 0.56, $Di^{F/F}$: 7.10 ± 0.37, $Ascl^{CreER}$;$Di^{F/F}$: 7.71 ± 0.34 for granule cells that are significantly tuned in both baseline and CNO injected sessions of the context discrimination task.

**E)** Fraction of cross-session registered cells found in Context A' that encoded space during exposure to same or different contexts in $Di^{F/F}$ controls and $Ascl^{CreER}$;$Di^{F/F}$ mice. $\chi^2$=2.79, p=0.42.

**F)** Fraction of cross-session registered cells found in Conext A' that did not encode space during exposure to same or different contexts in $Di^{F/F}$ and $Ascl^{CreER}$;$Di^{F/F}$ mice. $\chi^2$=1.27, p=0.73

**G)** Distribution of GCs' CNO induced normalized rate differences in $Ascl^{CreER}$;$Di^{F/F}$ mice compared to those in $Di^{F/F}$ controls. $P$ = 0.29, $Di^{F/F}$: 0.046 ± 0.017, $Ascl^{CreER}$;$Di^{F/F}$: 0.0413 ± 0.0172

**H)** The distribution of the context selectivity of spatially tuned cells that are disinhibited (right, **$P$ = 0.002) or inhibited (left, $P$=0.39) by CNO in $Ascl^{CreER}$;$Di^{F/F}$ mice compared to those in $Di^{F/F}$ controls.
Gray areas represent 2.5th and 97.5th% of null distributions for each cell type in B, C, E, F The null distributions are generated for each cell type by randomly permuting cell IDs of all cross-registered neurons (5,000 shufflings) and determining overlap among cell types. A-D,G,H: Two-sample Kolmogorov-Smirnov test. B-C-E-F: Kruskal Wallis test. Error bars, ± sem.

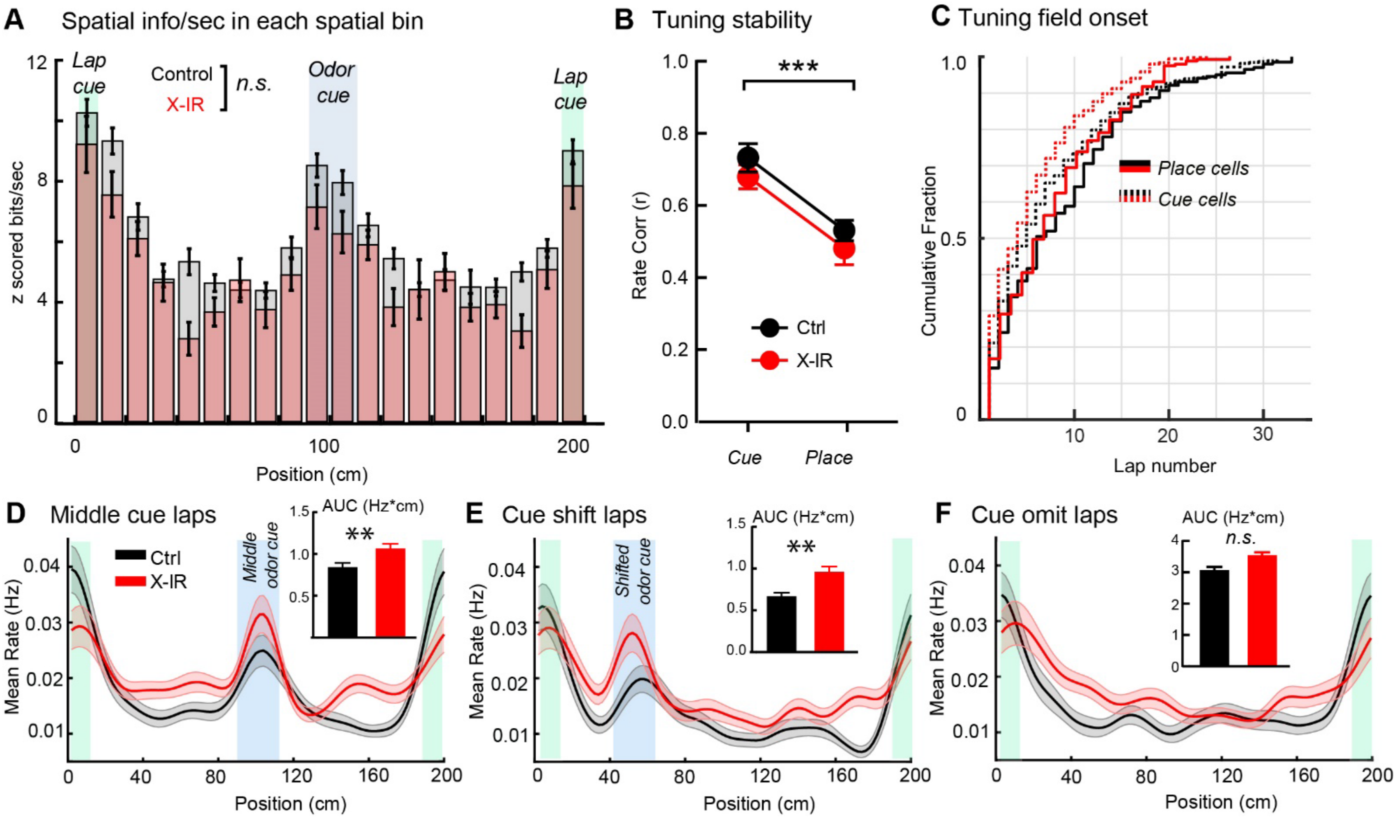


**Supplementary Figure 4. Effects of chronic ablation of neurogenesis on the functional segregation of cue and place responses in the DG (related to Figure 4).**

**A)** Mean spatial information by position showing similar enrichment of neurons with high spatial information around the sensory cues in both X-IR (pink) and control (gray) mice. Average Z scored spatial information for cells binned by tuning position during the spatial cue task performed by groups of mice. P=0.0926, Two-sample Kolmogorov-Smirnov test. Error bars, ± sem.

**B)** Tuning consistency of cue and place cells in control and X-IR mice during spatial cue task. Firing rate correlation between first and last halves of the session. Red dotted lines in violin plots show median, black dotted lines show quartiles. Interaction $F_{1,24}$ = 0.002, $P$ = 0.96; Cell Type $F_{1,24}$ = 29.12, ***$P$ < 0.0001; Genotype $F_{1,24}$ = 1.876, $P$=0.1834; 2-way ANOVA.

**C)** Emergence of cue and place responses. Cumulative distribution of spatial field onset lap for cue and place cells in control and X-IR mice during spatial cue task. Interaction $F_{1,617}$ = 0.006, $P$ = 0.94; Cell Type $F_{1,617}$ = 2.43, $P$ =0.12; Genotype $F_{1,617}$ = 6.004, *$P$=0.014; 2-way ANOVA.

**D)** Average spatial firing rates from all cells in Figure 4B during normal middle cue laps in control (black) and X-IR mice. (Inset) averaged area under the firing rate curves (AUC, Hz*cm) within the middle region corresponding to odor delivery during normal laps in control (black bar) and X-IR mice (red bar). Locations of the odor and lap cue are shown blue and green shaded areas, respectively. **$P_{Ctrl\text{-}IR}$ = 0.0036.

**E)** Average spatial firing rate by position for neurons on cue-shifted laps. (Inset) averaged area under the firing rate curves (AUC, Hz*cm) within the cue region during cue-shifted laps. **$P_{Ctrl\text{-}IR}$ =0.002.

**F)** Average spatial firing rate by position for neurons on cue-omitted laps. (Inset) averaged area under the firing rate curves (AUC, Hz*cm) during cue omitted laps. $P_{Ctrl\text{-}IR}$ =0.055. D-F: Mann-Whitney test. Error bars, ± sem.

**G)** Population vector (PV) correlations of granule cells with significant field at each treadmill position within a day in control (left) and X-IR (right) mice. For calculating PVs, lap-averaged spatial firing rate maps from normal middle cue laps of all tuned cells in Day1 were correlated with that of the second session from the dame day.

**H)** Population vector (PV) correlations of granule cells at each treadmill position within a week in control (left) and X-IR (right) mice. Lap-averaged spatial firing rate maps from normal middle cue laps of all tuned cells from the first session were correlated with that of the session imaged 7 days later.

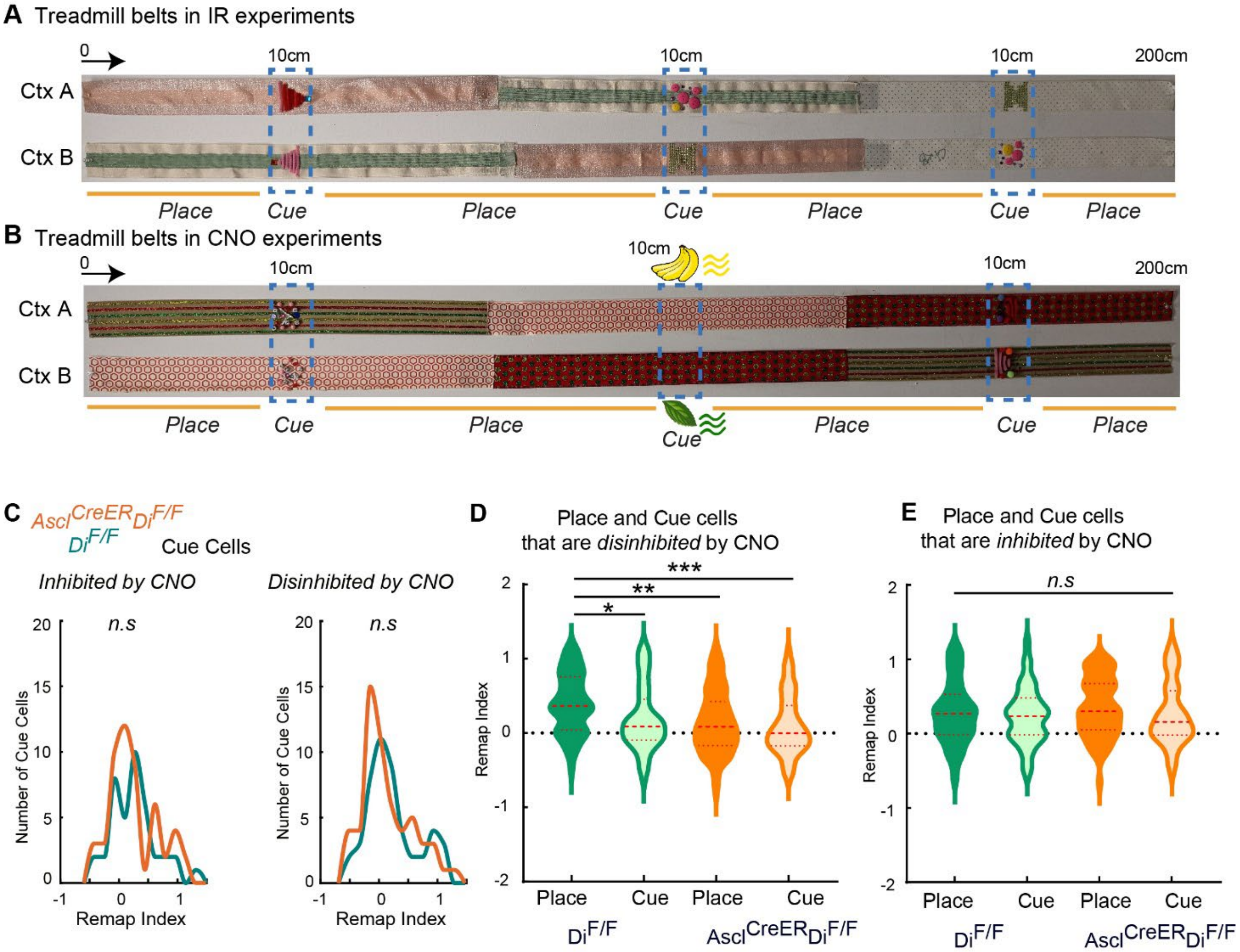


**Supplementary Figure 5: Effects of acute silencing of iGCs on the context selectivity of place and cue cells (related to Figure 5).**

**A)** Images of the two belts (A and B) used in the IR experiments. The order of the fabrics was changed and the three cues were affixed in a different arrangement. Cue encoding mGCs were defined as those with peak $Ca^{2+}$ event rate within the boundaries of the affixed tactile cues (10 cm wide) while the place encoding mGCs were defined as those with peak $Ca^{2+}$ event rate outside of the 10 cm boundary of the affixed tactile cues on the treadmill belt during the second exposure to context A. Direction of movement and the selected locations for the start of the belts are indicated.

**B)** Images of the two belts (A and B) used in the CNO experiments. The order of the fabrics was changed, different odor cues were presented at one location (1 sec pulse) and the two cues were affixed in a different arrangement. Cue encoding mGCs were defined as those with peak $Ca^{2+}$ event rate within the boundaries of the affixed tactile cues or the odor cue while the place encoding mGCs were defined as those with peak $Ca^{2+}$ event rate outside of the cue locations on the treadmill belt during the second exposure to context A. Direction of movement and the selected locations for the start of the belts are indicated.

**C)** The distribution of the remapping index of cue cells that are inhibited (left, $P$=0.63) or disinhibited (right, $P$ = 0.28 or by CNO in $Ascl^{CreER}$;$Di^{F/F}$ mice compared to those in $Di^{F/F}$ controls. Two-sample Kolmogorov-Smirnov test.

**D)** Comparisons of the remapping indices of place and cue cells that are disinhibited by CNO in $Di^{F/F}$ and $Ascl^{CreER}$;$Di^{F/F}$ mice. $\chi^2$=21.59, ***$P$ < 0.0001; *$P_{Cre\text{-} Place – Cre\text{-} Cue}$ = 0.039, **$P_{Cre\text{-} Place – Cre+ Place}$ = 0.0015, ***$P_{Cre\text{-} Place – Cre+ Cue}$ = 0.0002, Kruskal Wallis test, Dunn's multiple comparisons test. Dashed lines in violin plots show median, dotted lines show quartiles.

**E)** Comparisons of the remapping indices of place and cue cells that are inhibited by CNO in $Di^{F/F}$ and $Ascl^{CreER}$;$Di^{F/F}$ mice. $\chi^2$=2.7, $P$ = 0.43; Kruskal Wallis test.

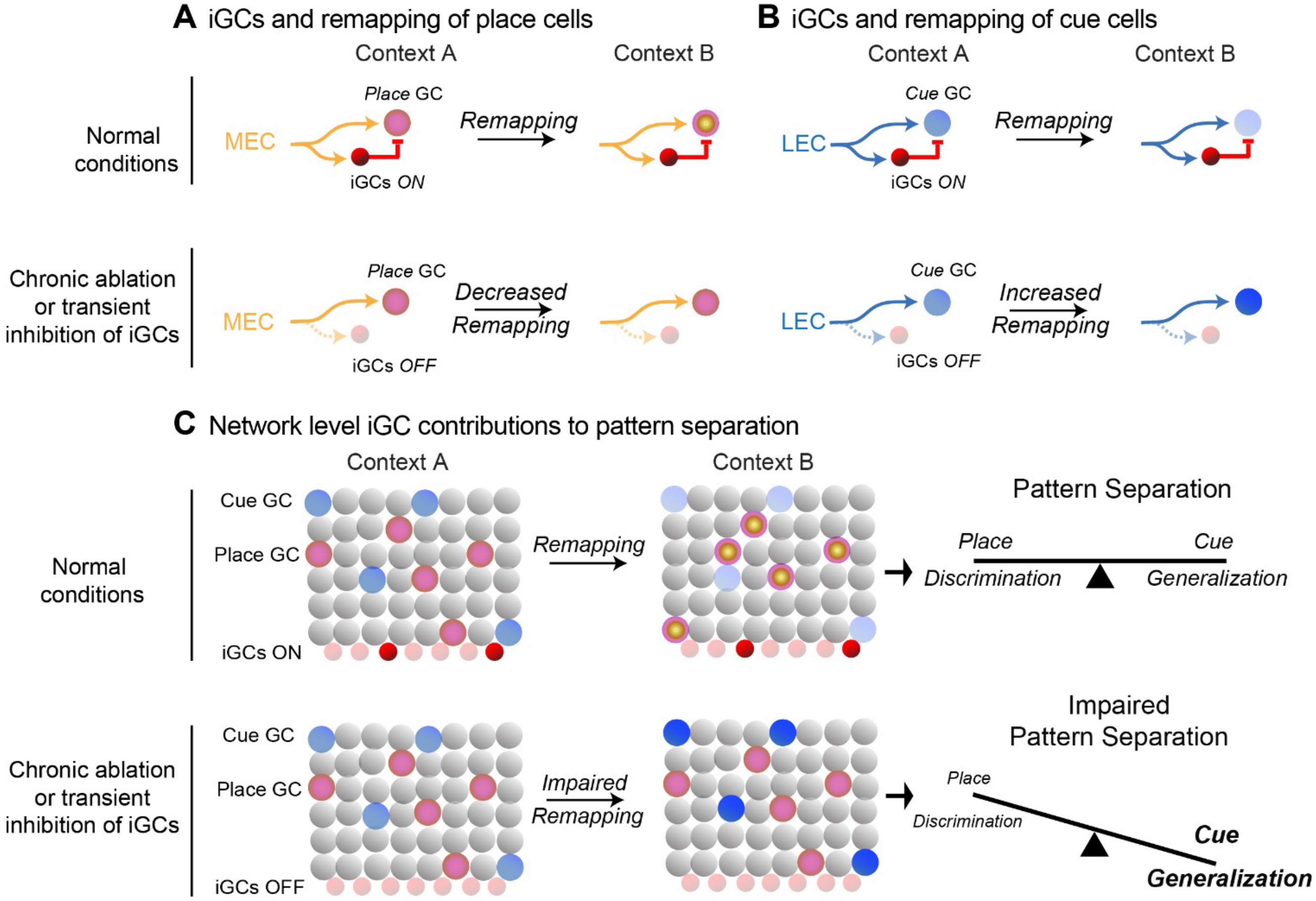


**Supplementary Figure 6: Model illustrating the contribution of adult born immature granule cells to pattern separation**

**A)** (Top) Under normal conditions, place encoding mature granule cells represent self-motion information between different cues context A (yellow circle with pink center) and represent self-motion information in context B by decorrelating their firing fields (yellow circle with pink surround). This is likely mediated by the modulation of spatial information arriving from the medial entorhinal cortex (MEC, orange arrows). (Bottom) Loss of iGCs prevent the modulation of these inputs to the place GCs, resulting in decreased remapping place GCs between context A and B (yellow circles with pink center).

**B)** (Top) Under normal conditions, cue encoding mature granule cells represent various sensory features of context A (light blue circle) and represent the same sensory cues in context B by modulating in their activity levels (lighter blue circle). This is likely mediated by the inhibitory modulation of sensory cue information arriving from the lateral entorhinal cortex (LEC, blue arrows) by immature adult born granule cells (iGCs, red circles). (Bottom) Chronic ablation of transient inhibition of iGCs prevent the inhibitory control of LEC inputs on cue GCs, resulting in high activity and increased remapping cue GCs between context A (light blue circle) and B (dark blue circle) even though the cues are at different locations of the sensory cues in different contexts.

**C)** Representations of contexts A and B are made up of mature GCs encoding sensory cues and place information. Under normal conditions, pattern separation of contexts A and B occurs by rate remapping of cue cells to form distinct traces for same objects encountered in different locations and global remapping of place cells to form non-overlapping representations about animal's location within different contexts. The balanced representation of sensory cue and self-motion information allows the network flexibly discriminate similar contexts and generalize common sensory features. Chronic ablation of transient inhibition of iGCs, impairs pattern separation by increasing remapping of sensory cue representations and decreasing the remapping of place cells and self-motion representations. In turn, this shifts the DG network state towards higher levels of sensory cue representation and less flexible self-motion representation resulting in over-generalization and lower discrimination.
